# Model-based fed-batch cultivation of *Viola odorata* plant cells exhibiting antimalarial and anticancer activity

**DOI:** 10.1101/2024.10.17.618783

**Authors:** R Babu, Manokaran Veeramani, Aadinath Wallepure, Vignesh Muthuvijayan, Shailja Singh, Smita Srivastava

**Author notes:** **Corresponding Authors: Smita Srivastava, Shailja Singh Email:** and. **The names, affiliations and e-mail addresses of co-authors are listed below:** Babu R^a^, Manokaran Veeramani^b^, Aadinath Wallepure^a^, Vignesh Muthuvijayan^a^. Department of Biotechnology, Bhupat and Jyoti Mehta School of Biosciences, Indian Institute of Technology Madras, Chennai, 600 036, India. Department of Chemical Engineering, Indian Institute of Technology Madras, Chennai, 600 036, India.

## Abstract

*Viola odorata* is used in the indigenous medicine to treat respiratory tract disorders and is limited in availability. Bioprocess principles can be applied to develop sustainable methods to produce high-quality *V. odorata* biomass. To this effect, a modified stirred tank reactor and a balloon-type bubble column reactor were used to improve biomass production. Nutrient feeding strategies were developed using first principle-based mathematical modelling to achieve higher cell density in the reactor. Experimental validation of the fed-batch model-predicted strategy resulted in a two-fold enhancement in biomass production (32.2 g DW L^-1^) at the bioreactor level. Also, bioreactor-cultivated biomass extracts were tested for *in vitro* hemolytic, cytotoxic, anti-inflammatory, and *in vivo* antiplasmodial activities. This is the first report on fed-batch cultivation in bioreactors and the antiplasmodial activity of *V. odorata*. Overall, the bioactive potential of the *in vitro*-generated biomass extracts is found to be similar to that in the natural plant biomass extracts.

**Highlights:**

- Cultivation of *V. odorata* cell suspension culture using a modified stirred tank and balloon-type bubble column bioreactors.
- Batch kinetic models were developed and extrapolated into a fed-batch model.
- Enhanced biomass production (32.2 g DW L^-1^) in bioreactors using a nutrient-feeding strategy.
- Extracts showed anti-inflammatory effects and up to 80 % inhibition of parasite growth, with no hemolytic activity.
- Confirmed antiplasmodial activity *in vivo*, effective alongside artesunate.
- *In vitro*-generated biomass extracts showed comparable bioactive potential to that of natural plants.

**Graphical Abstract:** 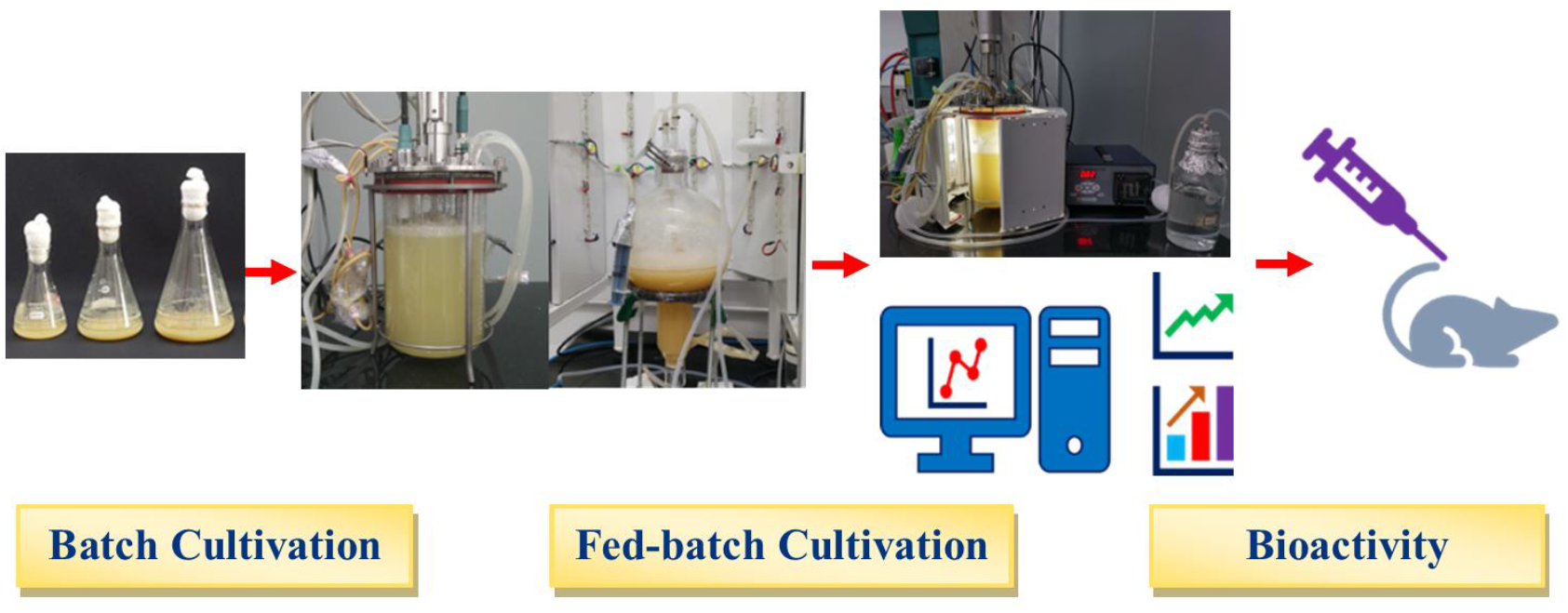

## 1. Introduction

Bioreactor-cultivated plant cell biomass can be an appropriate alternative to naturally sourced plant biomass from the wild. *In vitro* plant cell cultivation has produced high-value biomolecules and recombinant therapeutic molecules (Georgiev, 2014; Motolinía-Alcántara et al. 2021). The current study focuses on the production of herbal extracts from the bioreactor-cultivated plant biomass of the Indian medicinal plant *Viola odorata*, commonly called ‘Banafsha’. Banafsha is used in the ethnobotanical drug formulations comprised of leaves and flowers and is widely used in the indigenous system of medicine. *V. odorata* plant extract is used in combinations to treat fever, cold, cough, urinary disorders, kidney pain, and malaria (Singh, 1965). Due to various biological activities and commercial demand, the *V. odorata* plant is endangered in parts of India (Malik et al. 2011). *V. odorata* metabolite profile varies with the climatic and geographical conditions (Narayani et al. 2017b). Also, the commercial value of *V. odorata* plant biomass drives adulteration of other look-alike plants in *V. odorata’s* herbal raw materials (Singh, 1965). Unregulated uprooting has adverse effects on plant’s biodiversity. Also, microbial toxins and other contaminants reduce the quality of the raw materials collected from the wild. Hence, a sustainable *V. odorata* plant biomass production method is required to meet the commercial demand of *V. odorata* with appropriate herbal raw material quality as an alternative to natural plant biomass. For this, plant cell bioprocessing tools can be used to sustainably produce the bioactive principles from *V. odorata*. The protocols for the *in vitro* culture of *V. odorata* were reported by Narayani et al. (2017a). Further, the batch process condition of *V. odorata* cell suspension culture was established at the shake flask level (21.7 ± 0.8 g DW L^-1^) and 3 L stirred tank bioreactor (19.7 g DW L^-1^) with marine impeller (Babu and Srivastava, 2024). The reduction in the biomass concentration and cell viability (data not shown) in the bioreactors compared to shake flasks is presumably due to the shear environment in the bioreactors (Babu and Srivastava, 2024). Additionally, in the batch process, nutrient limitation and inhibition can lead to limited biomass production.

Mostly, the available bioreactor configurations in the market are designed to cultivate microbial strains. The cellular requirements of microbial cells and plant cells are different. A list of specific requirements, in terms of process parameters for plant cell culture have been discussed earlier (Georgiev et al. 2009; Andrews and Roberts 2017; Werner et al. 2017; Motolinía-Alcántara et al. 2021). In particular, plant cells in suspension are relatively shear sensitive and tend to grow as cell aggregates. Therefore, mass transfer is limited in high-cell-density cultivation. Modifying the bioreactor configuration and operational procedure to provide a low shear environment with better mass-transfer can support high biomass production in plant cell bioreactors (Prakash and Srivastava, 2007). Also, substrate limitation and inhibition are the constraints associated with batch process. Thus, to increase the biomass productivity, fed-batch cultivation strategies (continuous or intermittent feeding) have been employed in literature for cell/organ cultures of various species like *Azadirachta indica* (Prakash and Srivastava 2006; Palavalli et al. 2012), *Rubus chamaemorus* (Nohynek et al. 2014), *Capsicum chinense* (Kehie et al. 2016), *Oplopanax elatus* (Jin et al. 2020) and *Rhodiola sachalinensis* (Hao et al. 2022). However, no report is currently available on the fed-batch cultivation of *V. odorata* cell suspension culture in bioreactors.

Malaria is a life-threatening disease caused by the parasite *Plasmodium falciparum* and *Plasmodium vivax*. The infected female *Anopheles* mosquito transmits the parasites to humans. In 2020, over 241 million malarial cases and 627 000 malarial deaths were reported worldwide (World Health Organization (WHO), 2021). Multiple drug resistance in *Plasmodium* strains is an emerging issue in malaria management (Thu et al. 2017). Hence, the search for potential plant-based drug molecules is essential to control and manage the malaria. Nworu et al. (2017) reported cyclotide-rich extracts obtained from *Oldenlandia affinis* to inhibit *Plasmodium* proliferation. Therefore, the current study was planned to test *V. odorata* biomass extracts for their anti-malarial activity under *in vitro* and *in vivo* conditions. *V. odorata* plant biomass extracts were also tested for *in vitro* anti-inflammatory activity associated with the *P. falciparum* infection.

Cancer has been a leading cause of morbidity and mortality worldwide. Plant extracts rich in bioactive phytochemicals can prevent and treat cancer by targeting multiple pathways involved in cancer development (Fakhri et al. 2022). Plant-based remedies could be an alternative low-cost strategy to control and manage various cancer (Fakhri et al. 2022). The suitability of plant extracts as non-toxic drugs has been assessed using a hemolytic assay. The bioactive molecules present in the plant extract interact with the red blood cell membrane and release hemoglobin. *V. odorata* extracts contain a mixture of cyclotides that have been reported to exhibit both hemolytic and non-hemolytic activities. Hence, we investigated the anticancer and hemolytic activities of *V. odorata* extracts obtained from bioreactor-cultivated biomass.

The current study used batch kinetic data to develop a mathematical model describing growth and substrate utilization kinetics of *V. odorata* cell suspension culture. Further, the batch model was extrapolated to the fed-batch, and suitable nutrient-feeding strategies were developed *in silico* to enhance *V. odorata* biomass productivity (g DW L^-1^ d^-1^) with minimum hit and trial experiments at the reactor level. This is the first report to demonstrate the fed-batch cultivation of *V. odorata* cell suspension cultures in bioreactors. The bioreactor cultivated *V. odorata* plant cell biomass extract was analyzed for its cytotoxic, hemolytic, anti-inflammatory, and anti-malarial activities. Also, the key bioactive principles present in the biomass extract were identified using Gas Chromatography–Mass Spectrometry (GCMS) analysis. This study established an alternative route of producing *V. odorata* plant biomass, independent of nature.

## 2. Materials and Methods

### 2.1 Batch growth kinetics of *V. odorata* cell suspension culture

*V. odorata VOP-4* cells were previously established and maintained via periodic sub-culturing, as reported by Narayani et al. (2017a). Shake flask and bioreactor cultivation of *V. odorata* cell suspension culture was carried out with optimum growth conditions established earlier (Babu and Srivastava, 2024). *V. odorata* cell suspension was grown in optimized medium composition (sucrose: 45.6 g L^-1^, ammonium chloride: 0.5 g L^-1^, potassium nitrate: 2.1 g L^-1^, and other nutrients were at MS media composition) supplemented with 3 mg L^-1^ of 2, 4 - D. The initial pH was set at 5.4. The experiments were performed with an inoculum density of 7 g DW L^-1^. The shake flasks were then kept in a shaker operating at 85 rpm and 16/8 h light/dark (200 lx) conditions at 26.6 °C (Babu and Srivastava, 2024). The flasks were harvested (in triplicates) at 24 h intervals for 20 days to estimate the biomass (g DW L^-1^) produced and substrate utilized (g L^-1^).

### 2.2 Batch bioreactor cultivation of *V. odorata* plant cells

*V. odorata* plant cell suspension culture cultivated in a Stirred Tank Reactor (STR) with setric impeller and a balloon-type bubble column bioreactor (BTBCB). For this, the batch bioreactor cultivation was performed in a 3 L STR bioreactor (Applikon, The Netherlands) with the working volume of 2.4 L. A low shear setric impeller design reported earlier (Prakash and Srivastava, 2007) was adopted in this study to cultivate plant cells in a stirred tank reactor configuration. The setric impeller was manufactured at the central workshop of the Indian Institute of Technology Madras, Chennai. The impeller diameter (D_i_) to tank diameter (D_T_) ratio and impeller blade width (W_B_) to impeller diameter (D_i_) ratio were kept at 0.33. In four blade configurations, the blade length (LB) to impeller diameter (D_i_) ratio was kept at 1.1. The blade angle was kept at 60 ° (Fig. 1a) to provide axial and radial flow. The initial agitation speed was kept at 85 rpm, and the initial aeration rate was at 0.1 vvm.

**Fig 1.**
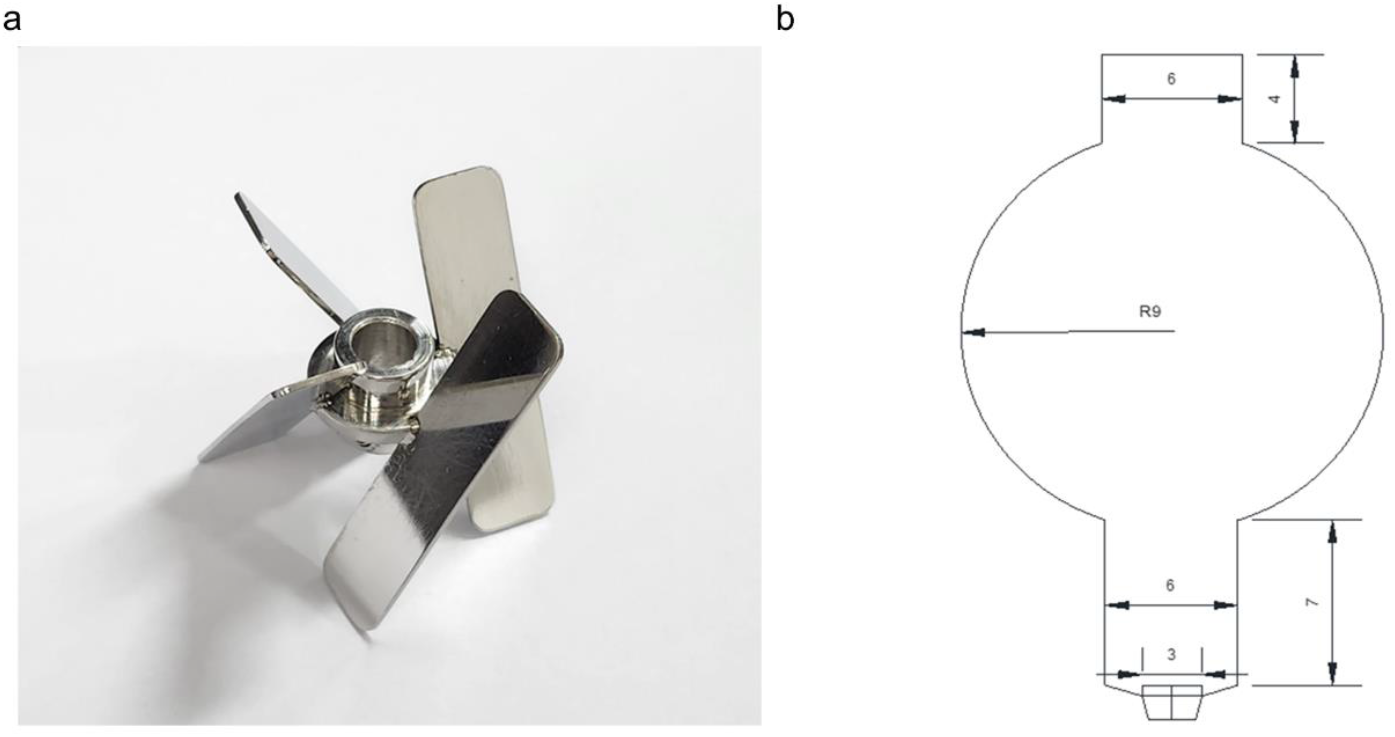
A setric impeller is used in a 3 L stirred tank reactor (a), and a balloon-type bubble column bioreactor (b) is used to cultivate a *V. odorata* cell suspension culture. The dimensions are mentioned in centimetres.

BTBCB was fabricated at the central glass blowing section of the Indian Institute of Technology Madras, Chennai. The dimension of the BTBCB is described in Fig. 1b. Grade four sintered glass sparger was fitted at the bottom of the vessel to aerate the culture broth with fine air bubbles. Silicone-based cork was fitted at the top to enable the air exit and enable a provision to collect the samples during cultivation. *V. odorata* plant cells were cultivated in the said BTBCB configuration with working volume of 1.5 L. The aeration was kept at an initial value of 0.2 vvm in the reactor. However, the agitation and aeration rates were varied during the cultivation period to prevent biomass from settling at the bottom of the bioreactors and to maintain dissolved oxygen >20 % during the cultivation period. The LED light jacket was used to maintain a 16/8 h light/dark photoperiod with 200 lx intensity, and the temperature was maintained at 26.6 °C by circulating cold/warm water through the cooling finger. *V. odorata* plant cells were cultivated in the optimized medium composition (described in section 2.1). The initial pH was adjusted to 5.4 before autoclaving and was not controlled during the cultivation. A silicone-based antifoam agent was used at a concentration of 0.012 % (v/v) to control the foaming (Babu and Srivastava, 2024). The samples were collected from the bioreactor at 24 h interval and analyzed for biomass (g DW L^-1^) and residual sucrose (g L^-1^) concentrations until 12 days of batch cultivation period.

### 2.3. Mathematical modelling assisted bioreactor cultivation of *V. odorata* plant cells

A maximum biomass concentration of 22.2 g DW L^-1^ was obtained under the optimised batch bioreactor cultivation conditions. *V. odorata* cell culture growth was limited by the nutrient availability in the batch cultivation. Hence, a fed-batch cultivation method was designed to overcome the substrate limitation/ inhibition and attain the high cell density in *V. odorata* cell suspension culture. To develop a kinetic model for subsequent process optimisation, basic knowledge of growth requirements and kinetic coefficients is needed for *V. odorata* cell suspension cultures. For this, *V. odorata* batch growth kinetics were obtained by cultivating it with optimised medium composition under optimal cultivation conditions (described in section 2.1).

#### 2.3.1 Inhibition of growth rate with respect to substrates carbon source and nitrogen source

The essential nutrients that limited *V. odorata* biomass production were identified earlier using Plackett-Burman and central composite design experiments (Babu and Srivastava, 2024). The experiments identified sucrose and potassium nitrate as the essential nutrients that positively affect biomass production. Therefore, it was decided to feed sucrose and potassium nitrate to the *V. odorata* cell suspension culture to enhance biomass production. Similarly, to avoid substrate inhibition during the fed-batch process, substrate inhibition was characterised using mathematical modelling. To this effect, *V. odorata* cell suspension culture was grown in an optimized medium composition supplemented with varying concentrations of sucrose (45, 75, 105, and 145 g L^-1^) and potassium nitrate (2, 4, 8, and 12 g L^-1^). The experiments were performed, and the flasks were harvested (in triplicates) at regular intervals of 2 days till 20 days to estimate the biomass (g DW L^-1^) produced, specific growth rate (μ; d^-1^) and substrate (sucrose and nitrate) utilized (g L^-1^).

#### 2.3.2 Development of a batch kinetic model for V. odorata cell suspension culture

A batch kinetic model was developed using the shake flask data. Further, a Luong-type inhibition model was incorporated to account for substrate inhibition due to nutrient feeding. Two different models (equations 1 and 2) and their combination (equation 3) were used to predict the growth of the cell culture in a batch.

(Model Type 1) Monod’s model (Srivastava and Srivastava, 2006) for growth limited by carbon and nitrogen

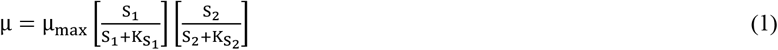

(Model Type 2) Sigmoid model (Prakash and Srivastava, 2006) for growth limited by carbon and nitrogen

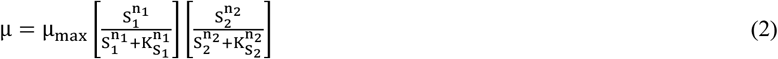

(Model Type 3) Monod’s and Sigmoid’s model (Prakash and Srivastava, 2006) combined to depict carbon and nitrogen-limited growth

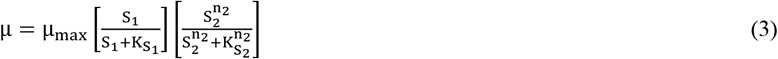

where,

μ = Specific growth rate in d^-1^

μ_max_ = Maximum specific growth rate with respect to limiting nutrients in d^-1^

S_1_ -Sucrose concentration in g L^-1^

S_2_ -Potassium nitrate concentration in g L^-1^

K_S1_ - Saturation concentration of sucrose in g L^-1^

K_S2_ - Saturation concentration of potassium nitrate in g L^-1^

The substrate inhibition by carbon and nitrogen was represented using the Luong-type model (Srivastava and Srivastava, 2006)

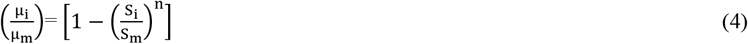

The assumptions made were as follows in the batch kinetic model.

1. Sucrose and potassium nitrate were the limiting nutrients affecting biomass production.
2. Higher concentrations of carbon and nitrogen sources inhibited biomass production.
3. Temperature and pH did not affect cell suspension growth in batch bioreactor cultivation.

Biomass accumulation and substrate consumption were represented with different model, including the following inhibition kinetics.

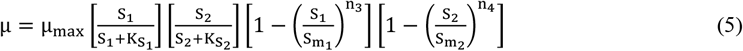

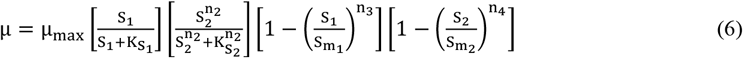

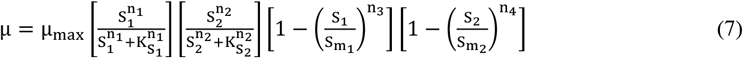

The batch kinetic data were used to predict the model parameters. The following equations represented the specific uptake rate of sucrose and potassium nitrate

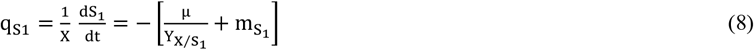

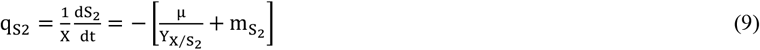

where,

q_S1_ = specific uptake rate of sucrose (S_1_)

q_S2_ = specific uptake rate of potassium nitrate (S_2_)

m_s1_ = maintenance coefficient based on sucrose

m_s2_ = maintenance coefficient based on potassium nitrate

Y _X/S1_ = yield of biomass with respect to sucrose

Y _X/S2_ = yield of biomass with respect to potassium nitrate

#### 2.3.3 Parameter estimation of batch kinetic models

*V. odorata* cells in suspension were cultivated under optimized conditions for twenty days. The samples were withdrawn every 24 h, and the batch growth kinetics were plotted based on the triplicate data. The experimental data were used to estimate the model parameters. The model parameters were optimized by the protocol reported by Srivastava and Srivastava (2008) wherein an iterative numerical technique was used to minimize the difference between the model predicted and experimental biomass data. Briefly, the computer software MATLAB^®^ (MathWorks, USA) was used to minimize the difference between the model prediction and experimental data. Runge-Kutta method of fourth order-based integration method was used for the model predictions from the growth kinetic equations. Rosenbrock optimization algorithm was used to find the minimum of a multivariable function. The following minimization criteria was used in the study.

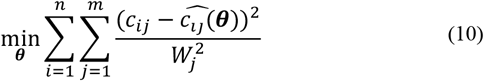

where,

*θ*^*L*^ ≤ *θ* ≤ *θ*^*U*^, c_ij_ – measured concentrations, 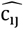-predicted concentrations, W_j_ – weights of each species, θ – parameters, θ^L^ - lower bound, θ^U^ - upper bound, n-number of data points, and m-number of species.

The model parameters were estimated using a non-linear regression technique (Prakash and Srivastava, 2006). The model prediction was compared with a separate set of experimental data to validate the developed batch model.

Parameter sensitivity analysis was performed to measure the change in the model predictions upon changes in the model parameters. If the changes in one parameter did not affect the model prediction, the parameter could be removed from the model, or a constant value can be assigned to it, which helps reduce the calculation and computation time. Absolute parameter sensitivity indicates the influence of individual parameters on the model predictions. It can be expressed as follows.

Absolute parameter sensitivity = 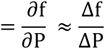

where,

f – optimized function – biomass concentration

P – parameters

Relative parameter sensitivity = 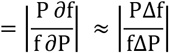

Further, absolute parameters were ranked using the relative parameter sensitivity based on their technological importance. A higher parameter value indicates a higher impact of the parameter on the process (Votruba et al. 1986; Goswami and Srivastava, 2000).

All three models (Equations 5, 6 and 7) were compared to select a suitable batch kinetic model for *V. odorata* cell suspension culture. The models were compared statistically using the protocol reported by Mannervik (1982) to select the most suitable model for the batch growth prediction. The following two approaches were used,

Based on variance:

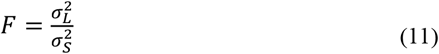

The variance calculated for the (Model Type 3) Monod’s and Sigmoid model was 0.093, and for the (Model Type 2) Sigmoid model was 0.097. The subscripts L and S represented the larger and smaller values of the variance of the developed model.

Based on objective function value:

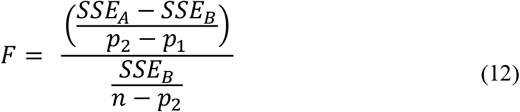

where,

SSE – Sum of squared errors values of Monod’s and Sigmoid model (SSE_A_) and Sigmoid model (SSE_B_)

n - number of data points used in the model development

pi - number of parameters present in the model (Monod’s and Sigmoid model (12) and Sigmoid model (13)) The F–test table values were calculated with 95 % confidence with appropriate degrees of freedom. The degrees of freedom for Monod’s and Sigmoid models was 22, and for the Sigmoid model was 21.

#### 2.3.4 Development of a fed-batch model for V. odorata cell suspension culture

The developed batch mathematical model was extrapolated to a fed-batch model by incorporating the effect of dilution terms to predict a suitable fed-batch cultivation strategy. The protocol reported in the literature by Prakash and Srivastava (2006) was adopted. The model equations (13-17) for the fed-batch process are shown below.

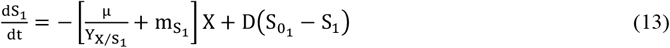

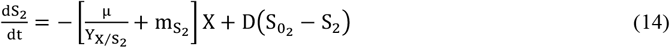

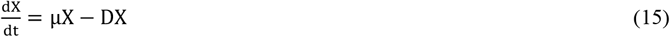

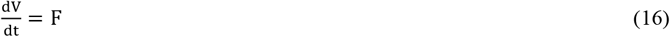

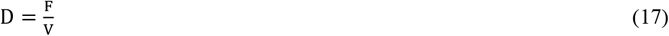

where,

F is the flowrate (L d^-1^), V is the volume (L), 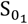 and 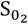 are initial sucrose and potassium nitrate concentrations (g L^-1^) in the feed bottle (fed-batch mode)

Fed-batch model equations 13 and 14 represented the accumulation rate of sucrose and potassium nitrate concentrations with respect to time. In addition, equations 16 and 17 represented the flow rate of the feed solution and dilution rate, respectively. The simulation was started as a batch and continued as a fed-batch, and respective model equations were simulated using MATLAB^®^ (MathWorks, USA) software to predict biomass and substrate concentration with respect to time.

The fed-batch process was designed using *in silico* strategies. The concentration of limiting nutrients (sucrose (45 to 400 g L^-1^) and potassium nitrate (2 to 30 g L^-1^)), volumetric flowrate (0.014 to 1.44 L d^-1^), initial medium volume in the reactor (0.8 to 1.5 L) and time of feeding (5 to 15^th^ day) were varied in a range *in silico* to predict growth and substrate kinetics. The MATLAB^®^ (MathWorks, USA) program computed the fed-batch process, and 29568 combinations were predicted. The results of the fed-batch process for different values of process parameters were simulated using model parameters adopted from the batch model. The process parameters combination leading to higher biomass productivity in comparison to batch, in minimum time was selected for experimental validation.

The fed-batch bioreactor cultivation of *V. odorata* plant cells was performed in both bioreactors, the STR and the BTBCB. Batch bioreactor cultivation conditions described in section 2.2 were employed for the fed-batch cultivation of *V. odorata* cell suspension culture. The initial working volume of the bioreactors was set at 1.2 L. Both bioreactors were allowed to run in batch mode up to the 10 days of the cultivation period, after which nutrients were continuously fed inside the reactor at a flow rate of 0.072 L d^-1^. The feed solution contained sucrose and potassium nitrate at concentrations were 250 g L^-1^ and 15 g L^-1^, respectively. The biomass was harvested (washed and filtered through filter paper under vacuum) after 17 days of cultivation to estimate the generated biomass concentration.

### 2.4 Analytical methods

The stigmasterol quantification protocol using HPLC was adapted from literature (Vemuri et al. 2018). The binary solvent system consisting of methanol and acetonitrile (95:5 %, v/v) in isocratic elution mode was used for separation and detection at a flow rate of 1 mL min^-1^. Thermo Scientific™ Hypersil GOLD™ HPLC column was used at 25 °C. The sample injection volume used was 20 μL with a PDA detector to measure the peak intensity at 208 nm. A standard plot was prepared in the 10 - 100 μg mL^-1^ concentration range.

The protocol reported by Xiao et al. (2013) was used for the quantification of phytol. Briefly, methanol and acetonitrile 30:70 (v/v) were used in isocratic elution mode with a 1 mL min^-1^ flow rate. A PDA detector was used at 210 nm to measure the peak intensity. Thermo Scientific™ Hypersil GOLD™ HPLC columns were used at 25 °C. The standard plot was made in the 10-200 μg mL^-1^ concentration range.

The powdered plant biomass extract was dissolved (10 mg mL ^-1^) in methanol and analyzed with the GCMS protocol reported earlier by Zhang et al. (2013). The HP-5ms Ultra Inert column (length 30 m, ID 0.25mm) was used. Electron ionization was done at 70 eV. The oven was operated isothermally at a temperature of 75 °C for 4 min. This was followed by a temperature rise of 5 °C min^-1^ up to 310 °C, held constant for 10 min. The sample injection temperature was 290 °C. The carrier gas flow rate (He) was 1.2 mL min^-1^, and the 1 μL sample was injected with a split ratio 1/15. The ion source and transfer line temperature were 230 °C and 280 °C, respectively. The mass spectra plot was acquired using full scan monitoring mode with a mass scan range of m/z 50−600. The solvent delay was kept at 3 min. Additionally, the protocol reported by Srikantan et al. (2018) was used for the residual sucrose, glucose, and fructose estimation.

### 2.5 Bioactivity analysis of *V. odorata* plant biomass extracts

For the study, the following *V. odorata* plant materials were used: D1 - natural plant biomass extract, D2 - *in vitro* plant biomass extract, D3 - somatic embryo biomass extract, D4 - stirrer tank reactor cultivated cell suspension biomass extract and D5 - airlift reactor cultivated cell suspension biomass extract (Babu and Srivastava, 2024). The biomass extracts were prepared using the protocol reported by Narayani et al. (2017a).

#### 2.5.1 Anticancer activity of V. odorata plant biomass extract

Caco-2 human colorectal adenocarcinoma cells and A549 – human lung adenocarcinoma cells were used to study the effect of plant extracts on the cancerous cell lines. Different concentrations (0.125 to 6 mg mL^-1^) of plant extracts (prepared as described in section 2.5) were tested on the cell lines, and the IC50 values were calculated using the protocol reported earlier by Mohinudeen et al. (2021). Briefly, the cell lines were maintained in a complete DMEM medium supplemented with 1 % (v/v) penicillin-streptomycin solution. Cells were maintained at 37 °C with 5 % (v/v) CO_2_. Cells were seeded in 96 well plates with a cell density of 1×10^4^ cells per well. Plant extracts were diluted in the medium to achieve different concentrations (0.125 to 6 mg mL^-1^) and added to each well on the plates. The plates were incubated for 48 h under the same conditions (37 °C with 5 % (v/v) CO_2_). The media was then discarded, and cells were washed multiple times with phosphate-buffered saline to remove traces of plant extracts. Then, alamarBlue solution (200 μL of 50 μg mL^-1^) was added to each well. The plates were incubated for 4 h at 37 °C. Cell viability was then measured using the absorbance at 570 and 600 nm wavelengths.

Cell viability was calculated as follows.

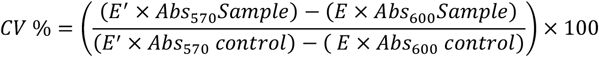

where,

*CV* = cell viability, *E* = molar extinction coefficient of oxidized alamarBlue at 570 nm, *E*′ = molar extinction coefficient of oxidized alamarBlue at 600 nm

#### 2.5.2 Hemolytic activity of V. odorata plant biomass extract

Hemolytic activity tests have been instrumental in identifying compounds with therapeutic potential, thereby guiding the development of novel drugs. All procedures involving human blood samples were conducted in accordance with protocols approved by the Institute Ethics Committee, IIT Madras (Approval No. IEC/2022-03/SS/10). A hemolytic assay was performed as per the protocol reported by Narayani et al. (2018) with certain modifications. In brief, human blood from a healthy volunteer was collected, and suspension of red blood cells (RBC) [1 % (v/v)] was prepared in phosphate-buffered saline (PBS). Further, 60 μL of the serially diluted plant extract (62.5–6000 μg mL^-1^ in PBS) sample and 60 μL of RBCs suspension were added to a 200 μL microcentrifuge tube and incubated at 37 °C for 1 h. As a test sample control, 60 μL of serially diluted plant extract was mixed with 60 μL of PBS. Samples in the microcentrifuge tubes were centrifuged at 1500 rpm for 5 minutes. The absorbance of the supernatant was determined at 415 nm.

Percentage hemolysis was calculated as follows.

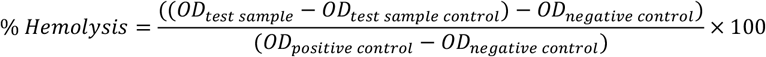

#### 2.5.3 Anti-inflammatory activity of V. odorata plant biomass extract

The anti-inflammatory activity of *V. odorata* plant extracts was studied on murine macrophage RAW 264.7 cell line. Before the study, the macrophages were maintained in RPMI 1640 medium supplemented with 10 % fetal bovine serum, 100 U L^-1^ of penicillin, and streptomycin. The cultures were incubated in a humidified atmosphere at 37 °C with 5 % CO_2_. The stock solution of plant extracts were prepared in phosphate buffer saline (pH 7.4) and diluted to 150 μg mL^-1^. The macrophages were subjected to different treatments: plant extracts in the presence of *P. falciparum*, plant extracts alone, PBS (negative control), lipopolysaccharides (LPS) (positive control), and *P. falciparum* alone. The treatments were applied for 4 hours. The cultures were incubated for 24 hours, and the culture broth was collected for the cytokine analysis.

Quantitative reverse transcription polymerase chain reaction (qRT-PCR) was done for gene expression analysis using the protocol reported by Vergallo et al. (2013), Bakir et al. (2011), and Jennings et al. (1997). Total RNA from *P. falciparum*-infected macrophage was isolated at appropriate time point (24 h) using the TRIZOL reagent (Invitrogen, Grand Island, NY) and quantified using a Nanodrop ND-1000 spectrophotometer (Thermo Fischer, USA). cDNA was prepared from 1 μg of RNase-free DNase-treated total RNA using first-strand cDNA Synthesis Kit (Thermo Fischer Scientific, USA), as per manufacturer’s instructions, using random hexamer primers. PCR reactions were conducted in Applied Biosystems, Real-Time PCR System (ABI, CA, USA) using PowerUp SYBR Green PCR Master Mix (Thermo Fisher Scientific, USA). The details of the primers (sequences and annealing temperatures) used were: TNFα: Forward 5’- CCTCTCTCTAATCAGCCCTCTG-3’, Reverse 5’ - GAGGACCTGGGAGTAGATGAG-3’ (T_a_- 60 °C); IL-10: Forward 5’ –TTAAGGGTTACCTGGGTTGC -3’, Reverse 5’-TGAGGGTCTTCAGGTTCTCC -3’ (T_a_- 55 °C); IL-12: Forward 5’ – ATGCCCCTGGAGAAATGGTG-3’, Reverse 5’ –GGCCAGCATCTCCAAACTCT -3’ (T_a_- 55 °C); IL-32ϒ: Forward 5’-AGGCCCGAATGGTAATGCT-3’; Reverse 5’-CCACAGTGTCCTCAGTGTCACA-3’ (T_a_- 60 °C). The thermal profile for the real-time PCR was as follows: amplification at 50 °C for 2 min, followed by 40 cycles at 95 °C for 15 sec, 60 °C for 30 sec, and 72 °C for 1 min. Melting curves were generated along with the mean C_T_ values and confirmed the generation of a specific PCR product. As an internal control for normalization, amplification of RNU6AP (RNA, U6 small nuclear 1; THP-1 cells) was used. The results were expressed as fold change of uninfected samples (control) using the 2-ΔΔCT method. The amplification of RNU6AP was conducted using specific primers: forward primer 5’-GGCCCAGCAGTACCTGTTTA-3’ and reverse primer 5’-AGATGGCGGAGGTGCAG-3’, with a recommended annealing temperature (T_a_) range of 55-60 °C.

#### 2.5.4 Anti-malarial activity of V. odorata plant extracts

##### In vitro growth inhibition assay

*Plasmodium falciparum* 3D7 (chloroquine (CQ) sensitive strain), RKL-9 (chloroquine-resistant strain), and R539T (artemisinin-resistant strain) strains (obtained from Malaria Research and Reference Reagent Resource Center - MR4) were cultured using the protocol reported by Sharma et al. (2021). *P. falciparum* strains were grown in human O^+^ RBCs along with RPMI1640 medium. The growth medium contains 0.2 % NaHCO_3_, 0.5 % (w/v) Albu Max II, 0.1 mM hypoxanthine, and 10 mg L^-1^ gentamicin. The cultures were incubated at 37 °C with a gas composition of 5 % O_2_, 5 % CO_2_, and 90 % N_2_ (v/v). Parasite culture was checked by scoring Giemsa-stained blood smears under a light microscope. Plant extract (250 μg mL^-1^) and vehicle control were incubated with parasites (at 1 % parasitemia with 2 % hematocrit) in the flat bottom 96 well plates, and growth inhibition was calculated as follows:

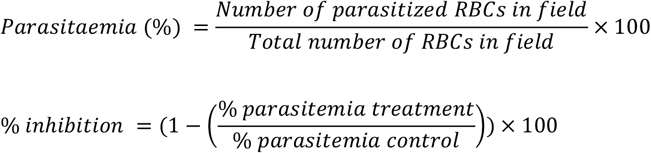

##### In vivo study

*Plasmodium berghei* was obtained from Malaria Research and Reference Reagent Resource Center (MR4) and maintained by continuous re-infection in mice. The animal experiments were conducted following the standard protocols approved by the Institutional Animal Ethics Committee, Jawaharlal Nehru University (IAEC No: 35/2019). The mouse blood was diluted with sterile saline to obtain 5 × 10^7^ parasitized RBC/mL. Infected blood (0.2 mL) containing 10^7^ parasitized RBC was inoculated into mice via IP using a 25G needle. Before administration, both the recipient and the donor mice were checked for parasitemia (microscopic examination of blood smear stained with Giemsa stain). Swiss albino mice (18 numbers - bred in-house at the animal facility of Jawaharlal Nehru University) were inoculated with 0.2 mL of infected blood containing 10^7^ parasitized RBCs and were assigned into six groups (n=3). One group was treated with saline (10 mL kg^-1^ d^-1^) as the negative control, and another group was treated with the standard anti-malarial drug Artesunate (ARTS:1.8 mg kg^-1^ d^-1^) as positive control for comparative analysis. The plant extract was administrated at 400 mg kg^-1^ d^-1^, and the plant extract was treated in combination with the standard anti-malarial drug ARTS (1.8 mg kg^-1^ d^-1^). All treatments started from day zero and lasted for four days. On the 4^th^ day, a thin blood smear was prepared and examined under a light microscope with Giemsa stain. The number of parasitized RBCs (using different fields) was calculated, and parasitemia (%) and inhibition of parasitemia (%) was estimated as given below.

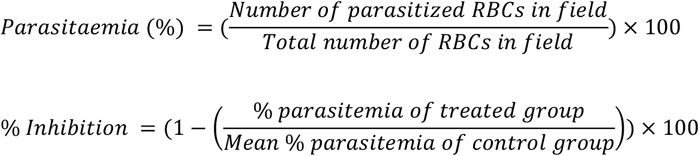

The Evans blue permeability test was used as previously described by Leten et al. (2014) and Uddin et al. (2022) to quantify vascular permeability in mice brains due to an increased parasite load. In brief, seven days after *P. berghei* infection, mice received an injection of 100 μL of 2 % Evans blue into the lateral tail vein. Brains were removed to evaluate the effects of Evans blue dye impregnation. Visual examination was done to compare the brains of the control and treated mice.

### 2.6 Statistical analysis

All the studies were done in replicates for reproducibility check and the data were represented as mean ± standard deviation (SD). The calculated means from OFAT experiments were analysed in a one-way analysis of variance (ANOVA) followed by Dunnett’s test for multiple comparisons. The mean difference was considered statistically significant when p < 0.05. Statistical analysis was performed using GraphPad Prism (v.9.5.1 - GraphPad Software Inc). Batch kinetic modelling and fed-batch cultivation strategies were developed using MATLAB^®^ software (MathWorks, USA).

## 3. Results and discussion

### 3.1 Batch growth kinetics of *V. odorata* cell suspension culture

*V. odorata* plant cell suspension culture was cultivated under the optimized culture conditions established earlier in the lab (Babu and Srivastava, 2024). The growth curve was plotted as shown in Fig. 2a. The maximum biomass concentration was obtained on day 12 with 21.7 ± 0.8 g DW L^-1^. The biomass productivity was calculated as 1.8 g DW L^-1^ d^-1^ with a biomass yield of 0.4 (g DW biomass produced per gram of sucrose utilized). The culture experiences a lag phase of 3 days and a log phase from 4 to 12 days. After that, the culture entered a stationary phase, due to nutrient limitation as shown in Fig. 2a.

**Fig 2.**
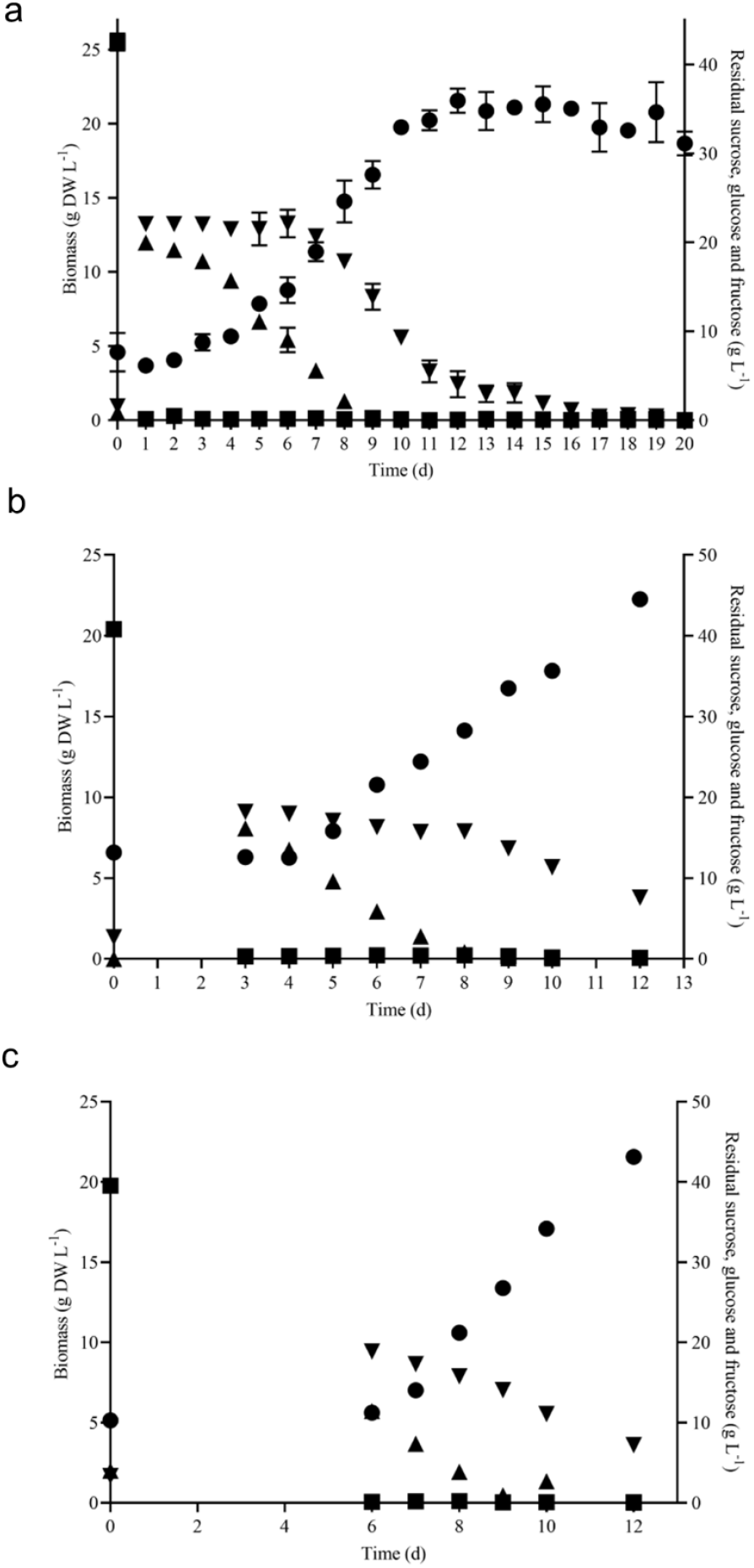
Batch growth kinetics of *V. odorata* cells on optimized medium in a shake flask (a), stirrer tank reactor with modified impeller (b) and balloon-type bubble column bioreactor (c). Biomass (*g DW L*^*-1*^, ●), sucrose (*g L*^*-1*^, ■), glucose (*g L*^*-1*^,▲) and fructose (*g L*^*-1*^,▼).

### 3.2 Batch bioreactor cultivation of *V. odorata* plant cells

The shake flask optimised culture conditions were implemented in the 3 L lab scale bioreactor to replicate the shake flask biomass productivity (1.8 g DW L^-1^ d^-1^) at the bioreactor scale. The bioreactor was configured with a marine-type impeller and sintered porous sparger to provide aeration and agitation. At the end of the batch cultivation (12 days), the maximum biomass concentration (19.7 g DW L^-1^) obtained was lower than the shake flask conditions (21.7 ± 0.8 g DW L^-1^) (Babu and Srivastava, 2024). Excessive foaming and a high-shear environment resulted in a lower biomass concentration. Hence, a low shear setric impeller was designed for STR to overcome the limitation associated with shear stress. Setric impeller provides both axial and radial mixing in the bioreactors. STR configured with a low shear setric impeller produced identical biomass concentration (22.2 g DW L^-1^) to shake flask conditions (21.7 ± 0.8 g DW L^-1^). Generated biomass (g DW L^-1^) and utilized substrate (g L^-1^) profile are presented in Fig. 2b. On the 8th day, glucose is completely consumed, and cells start utilising fructose (Fig. 2b) and 7.6 g L^-1^ was left in the medium on day 12. The electrical conductivity decreased from 3410 μS to 1323 μS on day 12.

A pneumatically driven BTBCB was designed based on the literature (Paek et al. 2001) to reproduce the shake flask biomass productivity. In the current study, the shake flask optimized culture conditions of *V. odorata* cell suspension culture were implemented, and a similar biomass concentration (21.6 ± 0.8 g DW L^-1^) was obtained with batch cultivation of 12 days. During the initial lag phase (0 - 4 days), excessive foaming was observed. Intermittent samples were withdrawn to estimate cell culture growth and substrate utilization kinetics (Fig. 2c). The spent medium analysis showed that sucrose and glucose were consumed entirely, and fructose up to 7.2 g L^-1^ still remained in the medium until day 12 of the batch cultivation period (Fig. 2c). The electrical conductivity decreased from 3570 μS to 1217 μS on day 12. BTBCB has been reported earlier for cultivating plant cells and organ culture (Paek et al. 2001). The concentric tube at the bottom and broader headspace provides better mixing with less foaming (Paek et al. 2001). BTBCB was used in the cultivation of *Panax ginseng* (Thanh et al. 2005), *Glycyrrhiza uralensis* (Wang et al. 2013), and carrot (Han et al. 2022) cell suspension for the production of ginsenoside, triterpenoid saponins, flavonoids and recombinant brazzein protein, respectively. Vigorous and intense foaming has been a limitation associated with aerated bioreactors. It was well documented for reducing biomass productivity via cell adhesion on the bioreactor walls and limiting the nutrient transfer to the cells. In BTBCB, the excessive foaming has been managed efficiently via the larger headspace available with this configuration. The concentric tube with a higher H: D ratio at the bottom of the BTBCB vessel supported the cultivation of plant cells at high cell densities. Sintered glass spargers provided fine air bubbles, resulting in a better oxygen transfer rate and mixing (Thanh et al. 2014).

### 3.3 Substrate inhibition studies

Sucrose and potassium nitrate were studied in a wide concentrations (described in section 2.3.1) range to find their inhibitory concentration. The growth curves were plotted on different substrate concentrations, and the concentration where the specific growth rate approached zero was noted as the complete inhibitory concentration for the respective nutrient. For sucrose (S_m1_) and potassium nitrate (S_m2_), the growth rate approaches zero around the concentration of 140 g L^-1^ and 12 g L^-1^, respectively (Fig. 3a and b). The data was used to estimate the dimensionless constant ‘n’ value in the model. The value of ‘n’ and respective inhibitory concentration estimated for sucrose and potassium nitrate from the model. The Monod and Sigmoid model combination predicted S_m1_ and S_m2_ as 179.24 g L^-1^ and 16.14 g L^-1^, respectively. Similar studies were conducted on *Azadirachta indica* cell suspension culture, and the inhibitory concentration of sucrose, potassium nitrate and di-hydrogen potassium phosphate were found to be 162.27 g L^-1^, 26.35 g L^-1^ and 0.63 g L^-1^, respectively (Prakash and Srivastava 2006).

**Fig 3.**
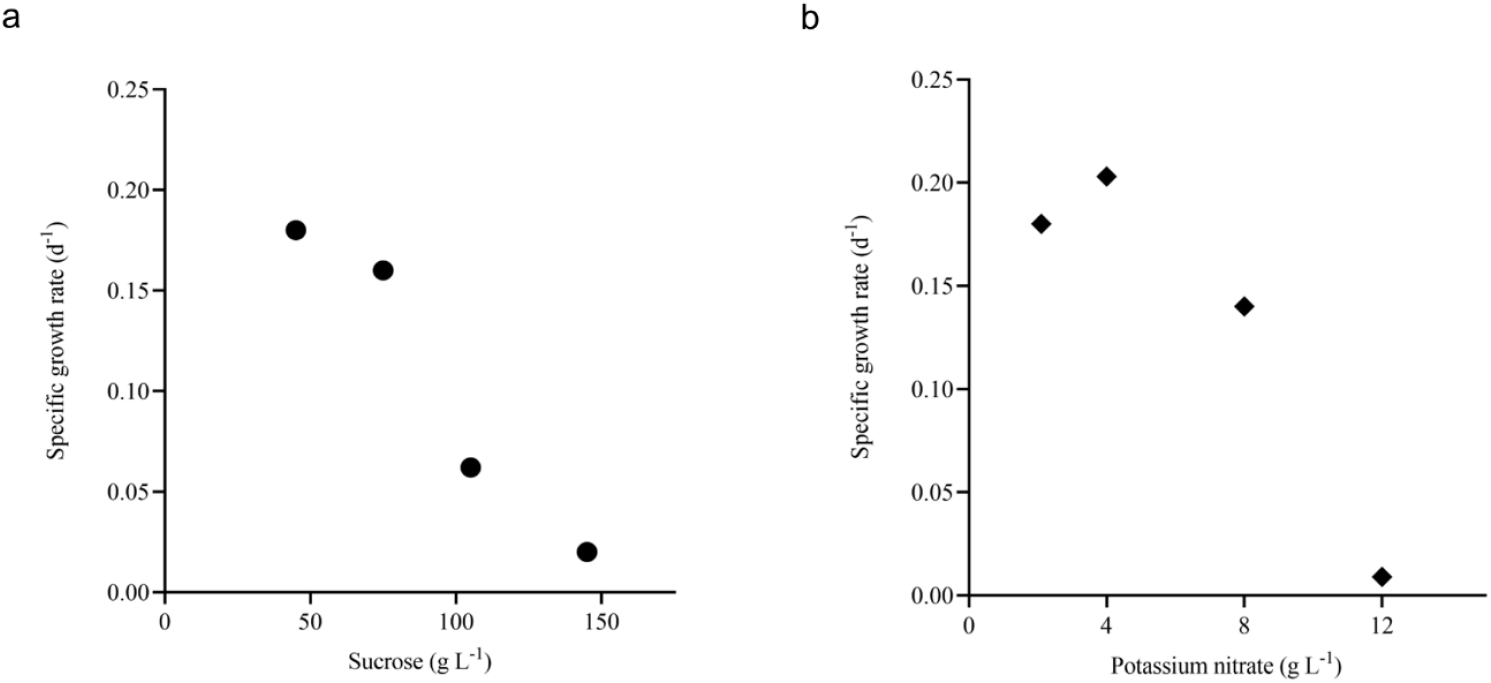
Inhibition of growth rate with respect to substrates. Plot (a) shows the specific growth rate with respect to different sucrose concentrations (*d*^*-1*^, ●) and (b) shows the specific growth rate with respect to different potassium nitrate concentrations (*d*^*-1*^, ♦).

### 3.4 Batch kinetic model for *V. odorata* cell suspension culture

The bounds for the parameter estimation were selected based on the experimental data and related works of literature (Table 1). The model parameters were estimated, and corresponding parameter values are presented in Table 2. Initially, Monod’s model was eliminated based on the value of S_m1._ The estimated value (85.93 g L^-1^) deviates from the experimental value (140 g L^-1^). Further, kinetic discrimination analysis was performed to select an optimal model with fewer model parameters, and the model parameter values are listed in Table 3.

**Table 1.** Parameter estimation: bounds for parameter estimation.

| Parameter | $Y_{X/S_1}$ | $Y_{X/S_2}$ | $\mu_{max}$ | $K_{S_1}$ | $K_{S_2}$ | $m_{S_1}$ | $m_{S_2}$ | $S_{m_1}$ | $S_{m_2}$ | $n_1$ | $n_2$ | $n_3$ | $n_4$ |
| --- | --- | --- | --- | --- | --- | --- | --- | --- | --- | --- | --- | --- | --- |
| $\theta^L$ | 0.1 | 0.1 | 0.1 | $1 \times 10^{-3}$ | $1 \times 10^{-3}$ | $1 \times 10^{-7}$ | $1 \times 10^{-7}$ | 44.9 | 2 | $1 \times 10^{-3}$ | $1 \times 10^{-3}$ | $1 \times 10^{-3}$ | $1 \times 10^{-3}$ |
| $\theta^U$ | 1 | 10 | 1 | 44.9 | 2 | 0.1 | 0.1 | 200 | 20 | 5 | 5 | 5 | 5 |

**Table 2.** Model parameter values for batch cultivation.

| Model | $Y_{X/S_1}$ | $Y_{X/S_2}$ | $\mu_{max}$ | $K_{S_1}$ | $K_{S_2}$ | $m_{S_1}$ | $m_{S_2}$ | $S_{m_1}$ | $S_{m_2}$ | $n_1$ | $n_2$ | $n_3$ | $n_4$ | Obj.fn value |
| --- | --- | --- | --- | --- | --- | --- | --- | --- | --- | --- | --- | --- | --- | --- |
| Monod | 0.51 | 7.86 | 0.40 | 29.93 | 0.17 | $5.08 \times 10^{-02}$ | $1.95 \times 10^{-05}$ | 85.93 | 3.67 | -- | -- | 3.68 | 2.16 | 2.068 |
| Monod and Sigmoid | 0.50 | 7.71 | 0.37 | 14.16 | 0.17 | 0.06 | $2.88 \times 10^{-04}$ | 179.24 | 16.14 | -- | 2.74 | 0.60 | 1.07 | 2.053 |
| Sigmoid | 0.47 | 7.76 | 0.37 | 13.51 | 0.01 | 0.04 | $2.34 \times 10^{-05}$ | 155.00 | 18.85 | 4.07 | 1.64 | 0.37 | 1.75 | 2.037 |

**Table 3.**
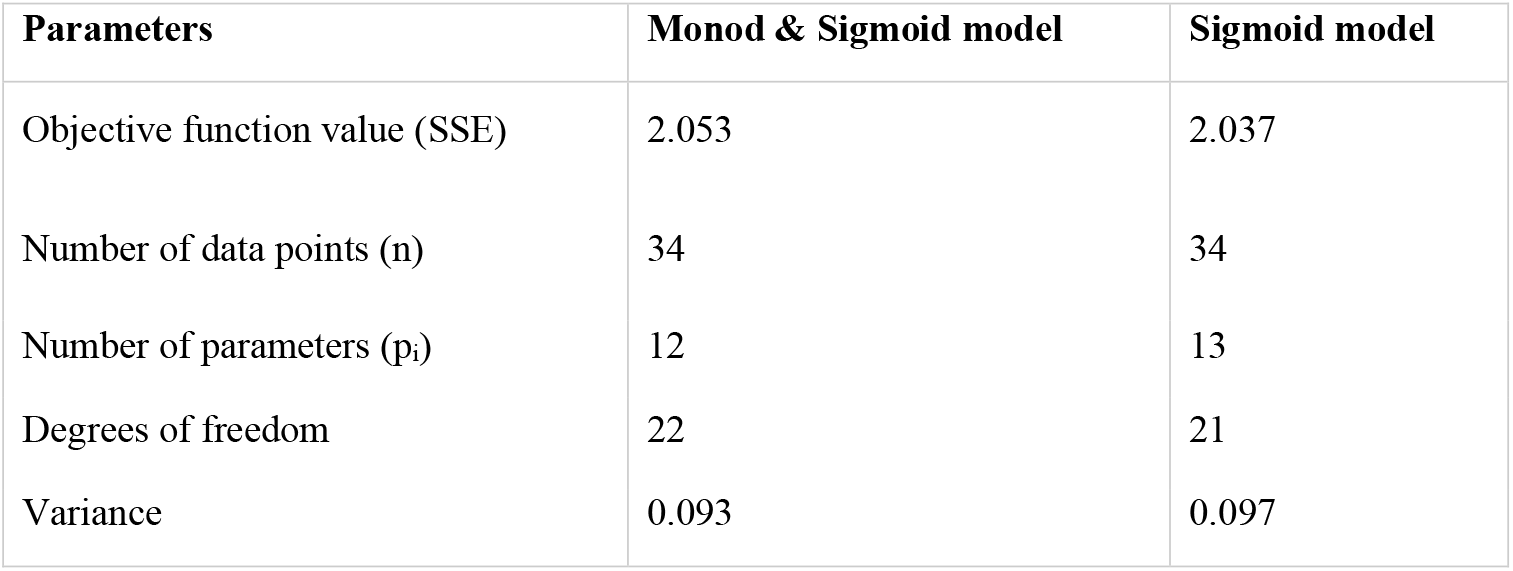
Kinetic model discrimination analysis.

Based on variance and Sum of Squared Errors (SSE), the Monod and Sigmoid model combination was compared with the Sigmoid model, resulting in the combination of both models being suitable for the prediction. Based on variance, the calculated F value (1.039) is lesser than the tabulated value (2.059) at a 95 % confidence interval and, based on the objective function value, the calculated F value (0.165) was lesser than the table value (4.325) at same confidence interval (95 %). Hence, the Monod and Sigmoid model combination was selected and analyzed for its parameter sensitivity. Based on the absolute parameter sensitivity, the parameter’s maximum growth rate and yield upon the potassium nitrate impacted the model response most significantly and were observed in the relative parameter sensitivity analysis (Table 4). The batch kinetic model was simulated using MATLAB^®^ (MathWorks, USA) software, and the prediction was validated with experimental data (Fig. 4a and b).

**Table 4.** Parameter sensitivity analysis of Monod’s and Sigmoid’s model (Model Type 3)

| Monod (sucrose) Sigmoid (potassium nitrate) model |  |  |
| --- | --- | --- |
| Parameter | APS | RPS |
| $Y_{X/S_1}$ | 0.12 | 0.03 |
| $Y_{X/S_2}$ | -0.06 | 0.21 |
| $\mu_{max}$ | -0.57 | 0.10 |
| $K_{S_1}$ | $7.43 \times 10^{-03}$ | 0.05 |
| $K_{S_2}$ | 0.25 | 0.02 |
| $m_{S_1}$ | -0.14 | $3.86 \times 10^{-03}$ |
| $m_{S_2}$ | 36.53 | 0.01 |
| $n_2$ | 0.02 | 0.02 |
| $S_{m_1}$ | $-1.18 \times 10^{-04}$ | 0.01 |
| $S_{m_2}$ | $2.42 \times 10^{-03}$ | 0.02 |
| $n_3$ | -0.17 | 0.05 |
| $n_4$ | $-7.43 \times 10^{-03}$ | $3.86 \times 10^{-03}$ |

APS - Absolute Parameter Sensitivity, RPS – Relative Parameter Sensitivity

**Fig 4.**
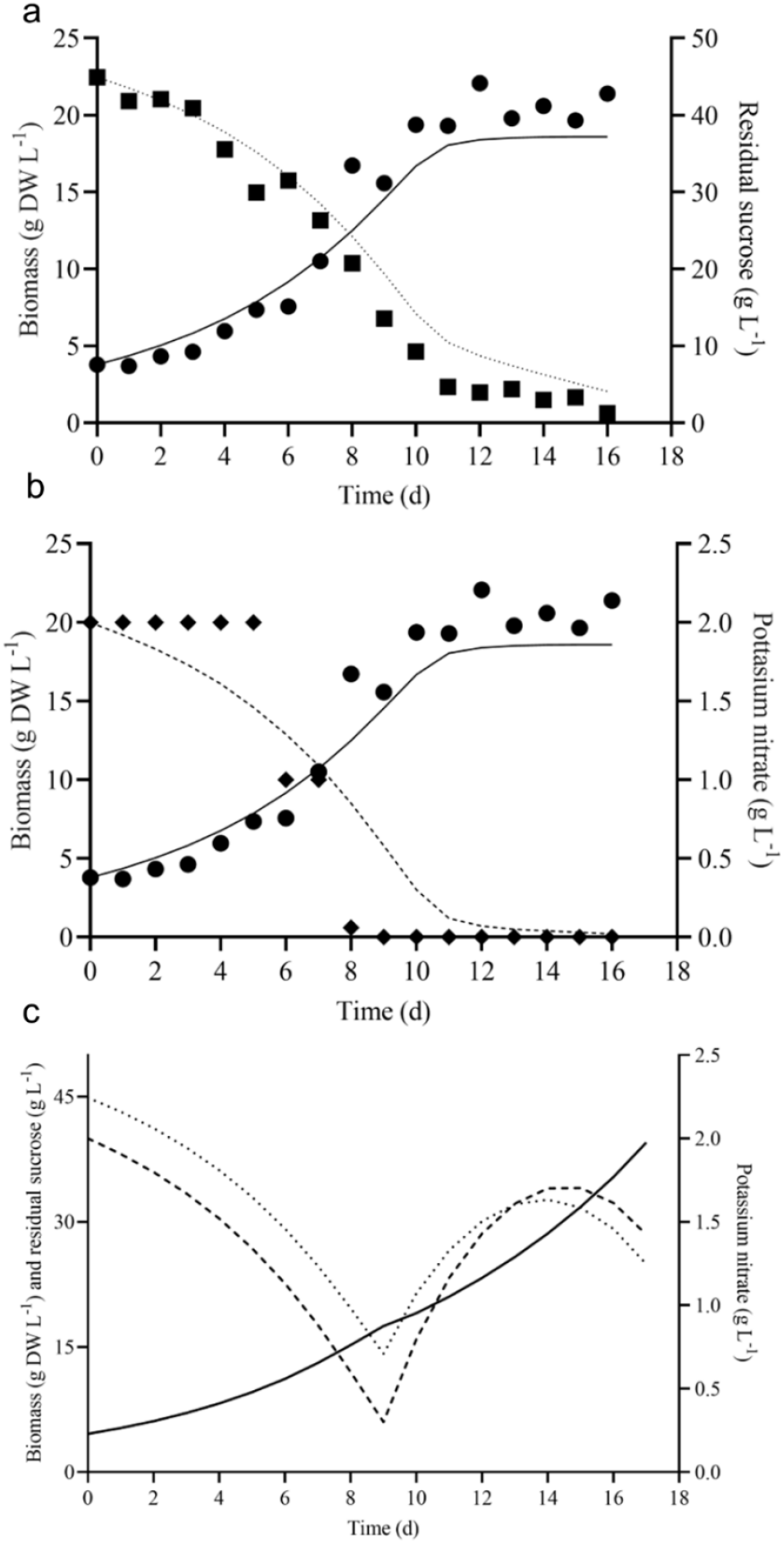
Model predicted and experimental batch growth kinetics of *V. odorata* cells on optimized medium. Plot (a) shows the experimental biomass production (*g DW L*^*-1*^, ●), sucrose (*g L*^*-1*^, ■) utilization and model predicted biomass production (*g DW L*^*-1*^, *continuous line*), sucrose (*g L*^*-1*^, *dotted line*) utilization; (b) shows the experimental biomass production (*g DW L*^*-1*^, ●), potassium nitrate (*g L*^*-1*^, ♦) utilization and model predicted biomass production (*g DW L*^*-1*^, *continuous line*), potassium nitrate (*g L*^*-1*^, *dashed line*). The fed-batch model predicted the growth kinetics of *V. odorata* cells on an optimized MS medium. Plot (c) shows the fed-batch model predicted biomass production (*g DW L*^*-1*^, *continuous line*), sucrose (*g L*^*-1*^, *dotted line*) utilization and potassium nitrate (*g L*^*-1*^, *dashed line*) utilization profile.

Batch kinetic models can be used to understand and optimize plant cell suspension cultures under *in-silico* conditions. The optimal cultivation conditions were obtained with *in-silico* predictions which were validated experimentally. The current study developed three growth models of *V. odorata* cell suspension culture (model type 1, 2 and 3), using batch growth and inhibition kinetic data. The batch model’s predictions were validated experimentally (Fig. 4a and b). Further, model-based fed-batch cultivation was carried out with the model predicted nutrient feeding strategy to achieve maximum biomass productivity (g DW L^-1^ d^-1^) in the *V. odorata* cell suspension culture.

### 3.5 Fed-batch cultivation of *V. odorata* plant cells in 3L bioreactors

Fed-batch cultivations are carried out to achieve maximum biomass productivity (g DW L^-1^ d^-1^) in plant cell suspension culture (Georgiev et al. 2009). The batch growth and substrate utilization of *V. odorata* cell suspension were modelled. The selected batch kinetic model was further extrapolated into a fed-batch model by incorporating the dilution terms (D). Various process parameters (described in section 2.3.4) were optimized by computer simulation using MATLAB^®^ software (MathWorks, USA). The feedstock solution concentrations of sucrose and potassium nitrate were 250 g L^-1^ and 15 g L^-1^, respectively. The reactor had an initial volume of 1.2 L. The fed-batch cultivation started on the 10^th^ day, and the feeding continued until the 17^th^ day with a flow rate of 0.072 L d^-1^. Under these conditions, the model-predicted biomass concentration was 39.52 g DW L^-1^, resulting in a model-predicted biomass productivity of 2.32 g DW L^-1^ d^-1^

The process conditions were selected based on the improved biomass productivity achieved in minimum time. The model predicted a suitable flow rate and feedstock solution concentration in such a way as to add nutrients into the *V. odorata* cell suspension culture and maintain it in a non-inhibitory concentration. *V. odorata* cell suspension was cultivated in batch mode for up to 10 days, and then the sucrose and potassium nitrate were fed at 0.072 L d^-1^. The model-predicted kinetic is represented in Fig. 4c. The experimental validation of the model prediction in a bioreactor (STR with two low shear setric impeller) obtained 32.2 g DW L^-1^ biomass with a productivity of 1.9 g DW L^-1^ d^-1^. The bioreactor was equipped with a setric impeller, and the limiting nutrients were fed to overcome the substrate limitations in the system. Presumably, the low shear setric impeller with an appropriate aeration rate provided a better hydrodynamic environment for the *V. odorata* cells’ growth. Modifications in the aeration and agitation system of the bioreactor were implemented in literature to maximize biomass production. For example, the setric impeller was used to cultivate the cell suspension culture of *Azadirachta indica* (Prakash and Srivastava 2006), and its hairy roots were cultivated with internal polyurethane foam support in an STR (Srivastava and Srivastava 2012). *Catharanthus roseus* (Jolicoeur et al. 1992) and *Eschscholtzia californica* (Archambault et al. 1994) were cultivated in bioreactors equipped with helical ribbon impellers. Recent advancements in bioreactor design and its limitations were discussed by Valdiani et al. (2019) and Motolinía-Alcántara et al. (2021).

BTBCB configuration was used in the large-scale cultivation of *Panax ginseng root* culture up to 500 L (Thanh et al. 2014). In the current study, *V. odorata* cell suspension culture was cultivated in batch and fed-batch mode using BTBCB. In the case of BTBCB, the fed-batch cultivation resulted in the biomass production of 27.4 g DW L^-1^ with a biomass productivity of 1.6 g DW L^-1^ d^-1^. BTBCB resulted in lower biomass productivity (1.6 g DW L^-1^ d^-1^) compared to batch cultivation. Biomass settling was observed toward the end (after – 10 days) of the fed-batch cultivation process. The airflow was adjusted in order to mitigate the settling of biomass at the base of the reactor. The biomass settled in the concentric tube at the bottom of the BTBCB which could be presumably responsible for the reduced biomass productivity.

The limited availability of medium nutrients can restrict the plant cell growth in a batch bioreactor. The growth phase of plant cells has been extended by feeding the limiting nutrients at a suitable concentration. Fed-batch cultivation of *Taxus chinensis* increased the biomass concentration to 27 g DW L^-1^ with taxane productivity of 9.3 mg L^-1^ d^-1^. The process used a low initial sucrose concentration (20 g L^-1^) and follows sucrose feeding (20 g L^-1^) on the 7^th^ day. The study used the single-factor approach, and at high concentrations of sucrose (> 30 g L^-1^), feeding resulted in osmotic stress to the cells (Wang et al. 1999). Hence, the model-based nutrient-feeding strategy was required to feed the limiting nutrients at non-toxic concentrations. Cultivation of *C. roseus* (Hyung et al. 1990), *A. indica* (Prakash and Srivastava 2006), and *Oplopanax elatus* (Jiang et al. 2021) have been reported using model-based fed-batch cultivation. In the current study, model-based fed-batch cultivation of *V. odorara* cell suspension culture resulted in a 3 - fold increase in biomass (32.2 g DW L^-1^) in the STR. However, mass transfer limitation due to high cell density and cell attachment on the bioreactor wall limited the growth of plant cells and the *V. odorata* biomass production in both the bioreactor configurations. Also, during the final cultivation phase, the dissolved oxygen concentrations experienced a decline and dropped below 20 %.

### 3.6 Quantification of stigmasterol and phytol in *V. odorata* plant biomass extracts

Stigmasterol is one of the most abundant plant sterols stabilizing the cell membrane structure and function. Kangsamaksin et al. (2017) reported that stigmasterol suppresses tumor growth via anti-angiogenic activity. Tumor cell proliferation has been inhibited by the downregulation of inflammatory cytokine (TNF-α) associated with the vascular endothelial growth factor pathway. *V. odorata in vitro* plant biomass extracts had the highest amount (206.1 μg g DW^-1^) of stigmasterol among other tested samples. Both bioreactors cultivated biomass extracts (D4 and D5) have a higher stigmasterol content than the natural plant source (45.5 μg g DW^-1^) (Fig. 5a). Also, stigmasterol content (0.007 μg mL^-1^) in natural plant leaf ethanolic extract was reported earlier (Aslam et al. 2020). Presumably, stigmasterol in *V. odorata* extracts can affect activity against the cancer cells (discussed in section 3.7.1).

**Fig 5.**
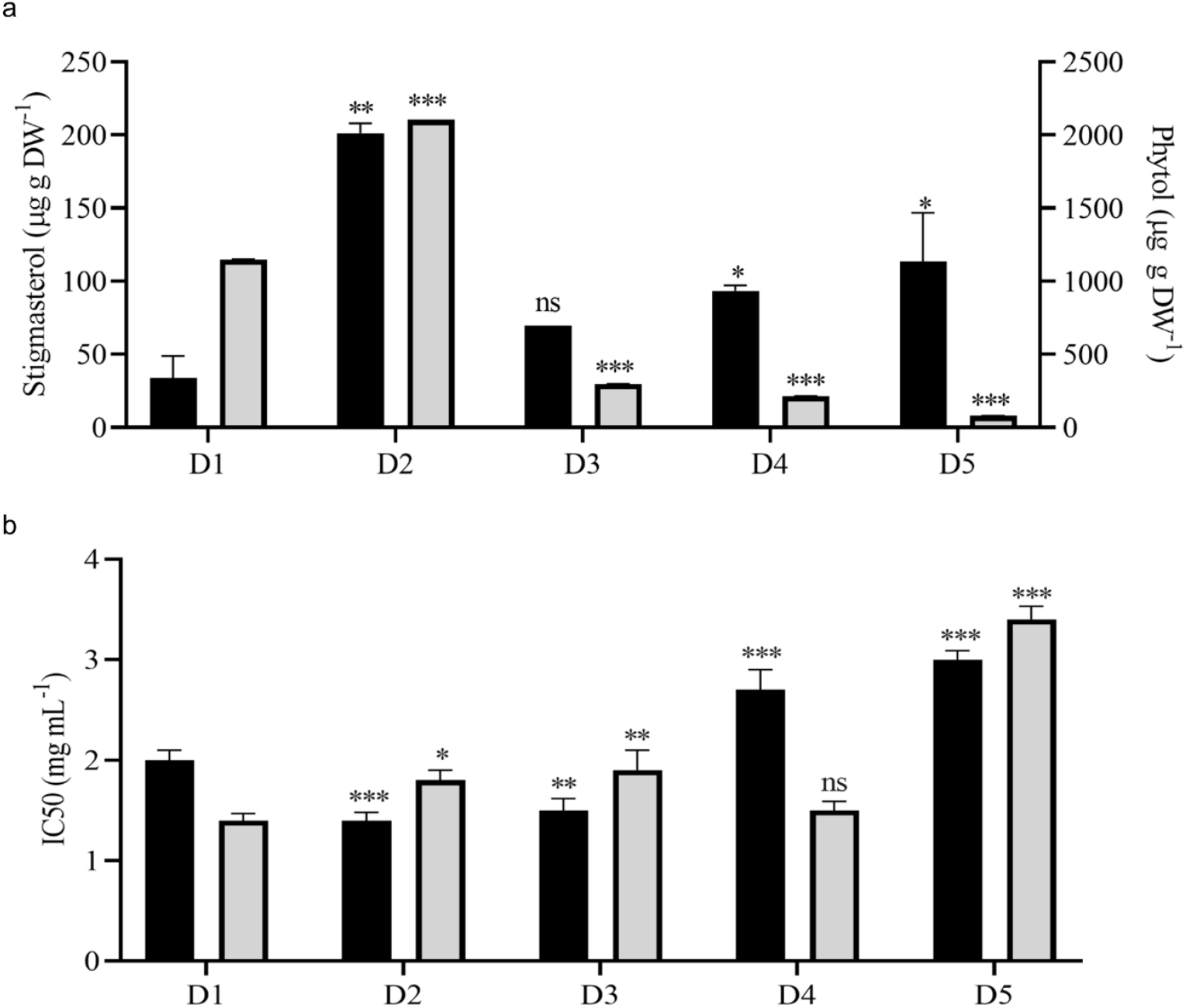
Metabolite quantification and cytotoxic activity of *V. odorata* plant biomass extracts. Plot (a) quantification of stigmasterol and phytol in *V. odorata* plant biomass extracts. Plot (b) cytotoxic activity of *V. odorata* plant extracts against A549 - Lung adenocarcinoma cells and Caco-2 human colorectal adenocarcinoma cells. D1 - natural plant biomass extract, D2 - *In vitro* plant biomass extract, D3 - somatic embryo biomass extract, D4 - stirrer tank reactor cultivated cell suspension biomass extract and D5 - airlift reactor cultivated cell suspension biomass extract. Different plant biomass extracts were compared with control (D1). *p <0.05, **p <0.01, and ***p <0.001. In (a), ■ – stigmasterol (μg g DW^-1^) and ■ – phytol (μg g DW^-1^). In (b), ■ – A549 - Lung adenocarcinoma cells and ■ – Caco-2 human colorectal adenocarcinoma cells.

Phytol is the long-chain unsaturated acyclic alcohol present in *V. odorata* biomass extracts. The *in vitro* plant biomass extract (D2) had the highest content (2102.9 μg g DW^-1^) of phytol among the tested samples (Fig. 5a), followed by natural plant extract (D1: 1149.4 μg g DW^-1^). STR-cultivated cell suspension biomass extract (D4) had a higher phytol content (214.3 μg g DW^-1^) than the airlift reactor cultivated cell suspension biomass extract (D5: 80.4 μg g DW^-1^). Phytol’s biological activities, including cytotoxic, anti-inflammatory, and antimicrobial, were discussed by Islam et al. (2018). Recently, Usman et al. (2021) reported that phytol (20 mg kg BW^-1^) reduces *P. berghei* parasitemia by about 25 % in Peter’s four-day suppressive test in the *in vivo* mice model. Earlier, phytol’s *in vitro* anti-malarial activity against *P. falciparum* was reported with an IC50 value of 18.9 μM (Grace et al. 2012). Primary and secondary metabolites in the *V. odorata* plant biomass extracts obtained with fed-batch cultivation were identified using GCMS (Fig. S1). A total of 23 metabolites were identified, including 5-hydroxymethylfurfural, 3-Deoxy-d-mannoic lactone, n-hexadecanoic acid and 4H-pyran-4-one, 2,3-dihydro-3,5-dihydroxy-6-methyl-which were the key metabolites present in the extract (Table S1).

In 1992, Cu et al. (1992) identified 23 volatile components in the *V. odorata* leaves using GCMS and GC-FTIR analysis. *V. odorata* leaves extract showed the presence of 25 essential oils, including butyl-2-ethylhexylphthalate and 5,6,7,7a-tetrahydro-4,4,7a-tri-methyl-2(4H)-benzofuranone as major constituents (Akhbari et al. 2012). *V. odorata* flower extract was analysed with GCMS and reported the presence of 84 compounds (Jasim et al. 2018). *V. odorata* leaf originating from France and Egypt was analysed, and 70 volatile compounds were reported (Saint-Lary et al. 2014). Plant parts and their geographical conditions, sample preparation, and extraction protocol impact the metabolite profile and the identification of phytochemicals from the plant biomass (Saint-Lary et al. 2014). In the current study, 23 key bioactive principles were identified through the GCMS analysis of plant extracts obtained with *in vitro* generated biomass obtained with fed batch STR cultivation. Along with cyclotides, other classes of phytochemicals such as flavonoids, phenolics, alkaloids, essential oils, saponins, tannins, anthocyanin, triterpenoids, and vitamin C were present in the *V. odorata* (Fazeenah and Quamri 2021). Also, suitable elicitation strategies can be developed to improve a specific metabolite content in the *V. odorata* cell suspension culture. On these lines, presence of specific bioactive compounds in this plant obtained by plant cell bioprocessing would be valuable in positioning their potential applications in medicine, addressed in the current study.

### 3.7 Bioactivity analysis of *V. odorata* plant biomass extracts

#### 3.7.1 Anticancer activity of V. odorata plant biomass extract

*V*.*odorata* plant extracts rich in cyclotides have been reported earlier to have cytotoxic activities (Narayani et al. 2018; Narayani et al. 2020). Cyclotides support the penetration of cancer drugs into the cancer cells by creating pores in the cancer cell membrane (Gould et al. 2012). In the current study, *in vitro*, organ culture extracts had similar bioactivity against A549 - lung adenocarcinoma and Caco-2 human colorectal adenocarcinoma cell lines (Fig. 5b). Bioreactor-cultivated biomass extracts have comparable activity (IC50: D4; 2.7 ± 0.2 mg mL^-1^ and D5; 3 ± 0.09 mg mL^-1^) against A549 - lung adenocarcinoma cells. In the case of Caco-2 human colorectal adenocarcinoma, STR cultivated biomass extract (IC50:D4; 1.5 ± 0.1 mg mL^-1^) had higher activity than airlift bioreactor extract (IC50: D5; 3.4 ± 0.13 mg mL^-1^). Overall, compared to the natural plant biomass extract (D1), the *in vitro-*generated biomass extract has comparable bioactivity against both cell lines (Fig. 5b). The cytotoxic activity of *V. odorata* plant biomass extracts (D1 to D5) was investigated on a non-cancerous L929 Fibroblast cell line. The results revealed a non-cytotoxic behaviour, indicating no adverse effects, even at a concentration of up to 6 mg mL^-1^ of the plant extract. This implies that the plant extract with cyclotides is specifically active against the cancer cells, and a similar observation was reported earlier (Narayani et al. 2018). *V. odorata* somatic embryo extracts exhibited specific cytotoxic effects against various cancer cell lines, including human colorectal adenocarcinoma Caco-2 cell line (IC50 = 370 μg mL^-1^), oral cancer UPCI: SCC131 cell line (IC50 = 93.60 μg mL^-1^), and cervical cancer Hela cell line (IC50 = 93.99 μg mL^-1^), as reported by Narayani et al. (2018). Notably, these extracts demonstrated non-cytotoxic behaviour toward the non-cancerous embryonic kidney cell line HEK 293 (IC50 = 426.1 μg ml^-1^). In another study, *V. odorata* hydro-alcoholic extract (5-200 μg mL^-1^) was studied in 4T1 breast cancer cells with MTT assay (1×10^4^ cells mL^-1^). *V. odorata* plant extract inhibited cancer cell growth at higher concentrations >100 ug mL^-1^ (Alipanah et al. 2018). Cyclotides were the key bioactive molecules in the *V. odorata* extracts and responsible for bioactive properties (Gerlach et al. 2010). The mechanism of action of cyclotide was membrane disruption (Svangård et al. 2007). In the current study, *in vitro*-generated plant biomass extracts were tested for cytotoxicity against A549 Lung adenocarcinoma cells and Caco-2 human colorectal adenocarcinoma cells. The biomass extracts inhibited the growth of both cancerous cells, with an IC50 value of less than ∼3 mg mL^-1^. These results indicate that *in vitro*-generated *V. odorata* biomass could be an alternative to natural plant biomass extracts for its cytotoxic applications.

#### 3.7.2 Hemolytic activity of V. odorata plant biomass extract

The present study focused on discovering plant extracts’ dose-dependent effect on hemolysis. At 1000 μg mL^-1^ concentration, all tested samples showed non-hemolytic behavior, and D1 and D2 started lysing the RBCs around the concentration greater than 1000 μg mL^-1^ (Fig. 6a). D3 showed only 5 % hemolysis at 2000 μg mL^-1^ (Fig. 6b). However, extracts D4 and D5 showed non-hemolytic behavior around the concentrations of 3500-4000 μg mL^-1^ (Fig. 6c and d). The non-hemolytic properties of *V. odorata* plant extracts indicate the suitability of the *in vitro* generated biomass extract as a therapeutic agent, and it could be used as a single or combination therapy along with other therapeutic molecules. This is also supported by its well-known ethnobotanical use in oral concoctions and wellness products. In the current study, the bioreactor-cultivated biomass extracts showed the non-hemolytic nature up to 4000 μg mL^-1^. In the literature, *V. odorata* somatic embryo extracts showed non-hemolytic activity at tested concentrations up to 1000 μg mL^-1^ (Narayani et al. 2018).

**Fig 6.**
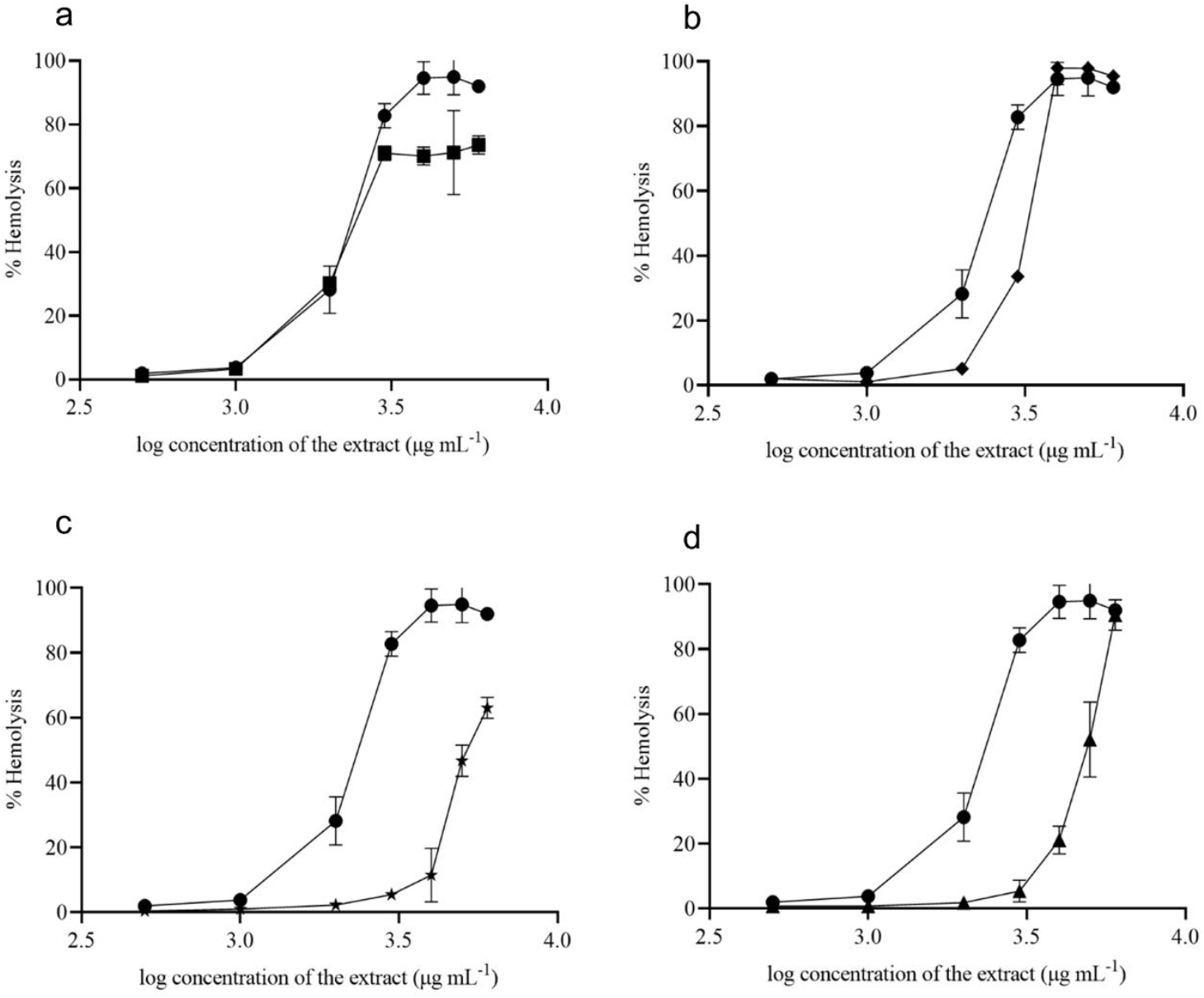
Hemolytic activity of *V. odorata* plant biomass extract. D1 - natural plant biomass extract (●), D2 - *In vitro* plant biomass extract (■), D3-somatic embryo biomass extract (♦), D4 -stirrer tank reactor cultivated cell suspension biomass extract (★) and D5 - Airlift reactor cultivated cell suspension biomass extract (▲). Plots (a-d) show % hemolysis of different plant biomass extracts compared with control (D1).

#### 3.7.3 Anti-inflammatory activity of V. odorata plant extracts

*In vitro*-produced *V. odorata* biomass extracts (D2 and D4 – selected based on their antiplasmodial activity) were tested with RAW 264.7 macrophage cell line. The macrophages were treated with plant extract, Lipopolysaccharide (LPS), and plant extract plus *P. falciparum*. Upon LPS stimulation, upregulation in inflammatory molecules was observed, and with *P. falciparum* infection, pro-inflammatory cytokines (TNF-α, IL12, IL32-ϒ) expression increases; however, anti-inflammatory cytokine (IL10) does not show any change. The pro-inflammatory cytokines were not upregulated when the macrophages were treated with the plant extracts, confirming the compatibility of *in vitro* plant biomass extracts. The plant extracts (D2 and D4) showed good inhibitory activity against inflammatory molecules. Upon *P. falciparum* infection, increased inflammation was observed in RAW 264.7 cells. However, adding plant extracts (D2 and D4) to the infected RAW 264.7 cells downregulate the inflammatory molecule expression (Fig. 7). The anti-inflammatory activity of *V. odorata* natural plant extracts in animal models was reported by Drozdova and Bubenchikov (2005) and Koochek et al. (2003).

**Fig 7.**
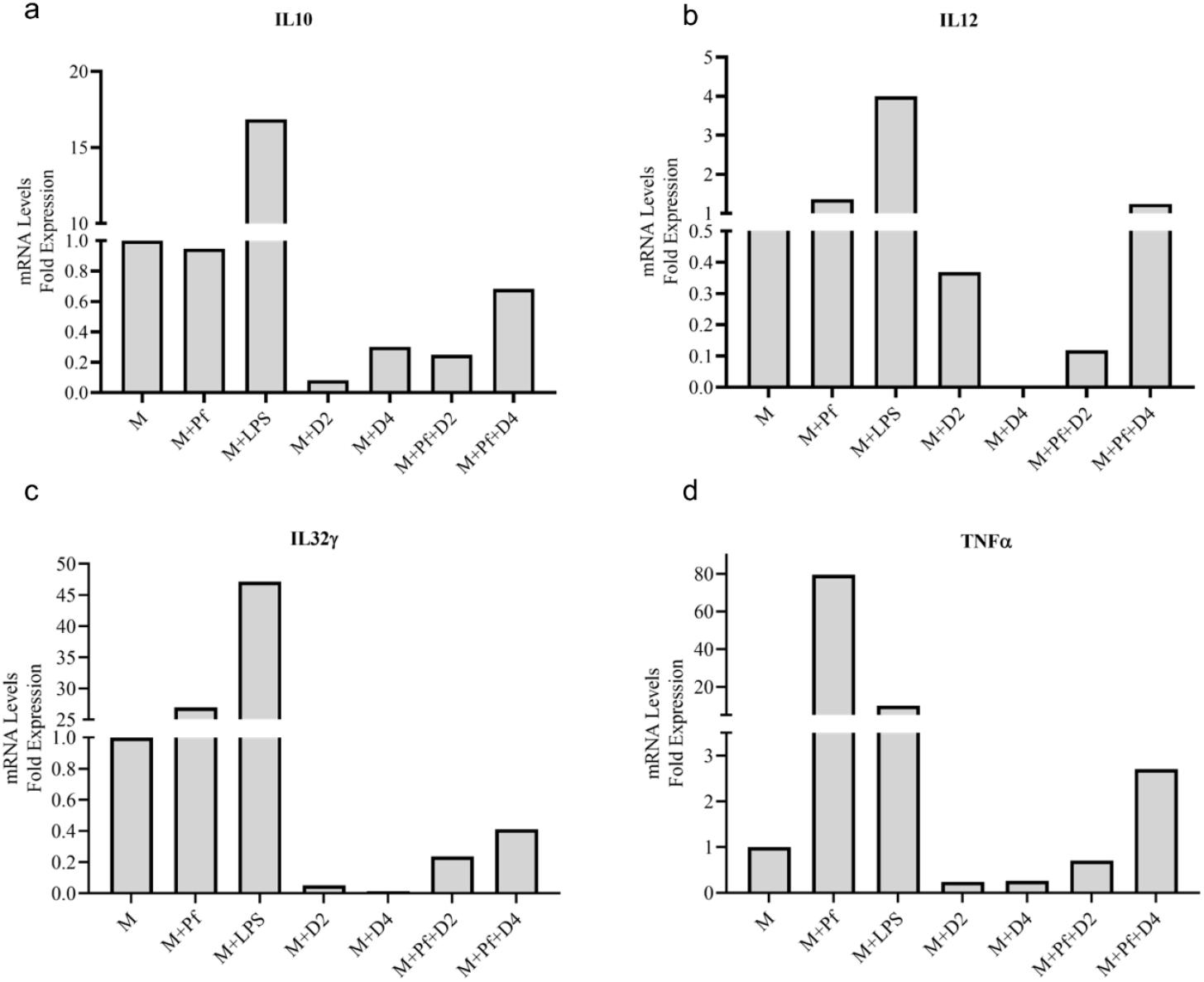
Anti-inflammatory activity of *V. odorata* plant extracts. Plots (a-d) show the level of mRNA expression of pro-inflammatory (TNF-α, IL12, IL32-) and anti-inflammatory cytokines (IL10). M - Macrophage RAW 264.7 cells, Pf *-P. falciparum*, LPS -Lipopolysaccharides, D2 - *In vitro* plant biomass extract and D4 -stirrer tank reactor cultivated cell suspension biomass extract.

*V. odorata* plant extracts containing secondary metabolites (Cu et al. 1992) and cyclotides (Ireland et al. 2006) were reported for their immunomodulatory activities (Dayani et al. 2022). Chronic inflammation has been associated with autoimmune disorders, cardiovascular diseases, neurodegenerative conditions, and specific cancers (Duan et al. 2019). Plant extracts with anti-inflammatory properties had the potential to mitigate and prevent diseases, as mentioned earlier (Oguntibeju 2018). Also, *V. odorata* plant extract has been used to treat inflammatory skin conditions, such as eczema or dermatitis (Batiha et al. 2023). *V*.*odorata* extracts often contain bioactive compounds, such as polyphenols, flavonoids, and alkaloids (Feyzabadi et al. 2017), that possess anti-inflammatory properties (Koochek et al. 2003). In the current study, *V. odorata* biomass extracts showed promising anti-inflammatory in the RAW 264.7 macrophage cell line (Fig. 7).

#### 3.7.4 The anti-plasmodial activity of V. odorata plant extracts

The antiplasmodial activity was tested in the three different *P. falciparum* strains (3D7, RKL9, and R539T). The natural and *in vitro* generated biomass extracts showed greater than 80 % inhibition activity against all three *P. falciparum* strains (Fig. 8). Extracts prepared from the biomass grown in the STR (D4) showed promising activity against the *P. falciparum* R539T strain and inhibited the parasite growth by more than 80 %. Further, extracts from the *in vitro* plants (D2) and biomass grown in the STR (D4) were studied in the *in vivo* models

**Fig 8.**
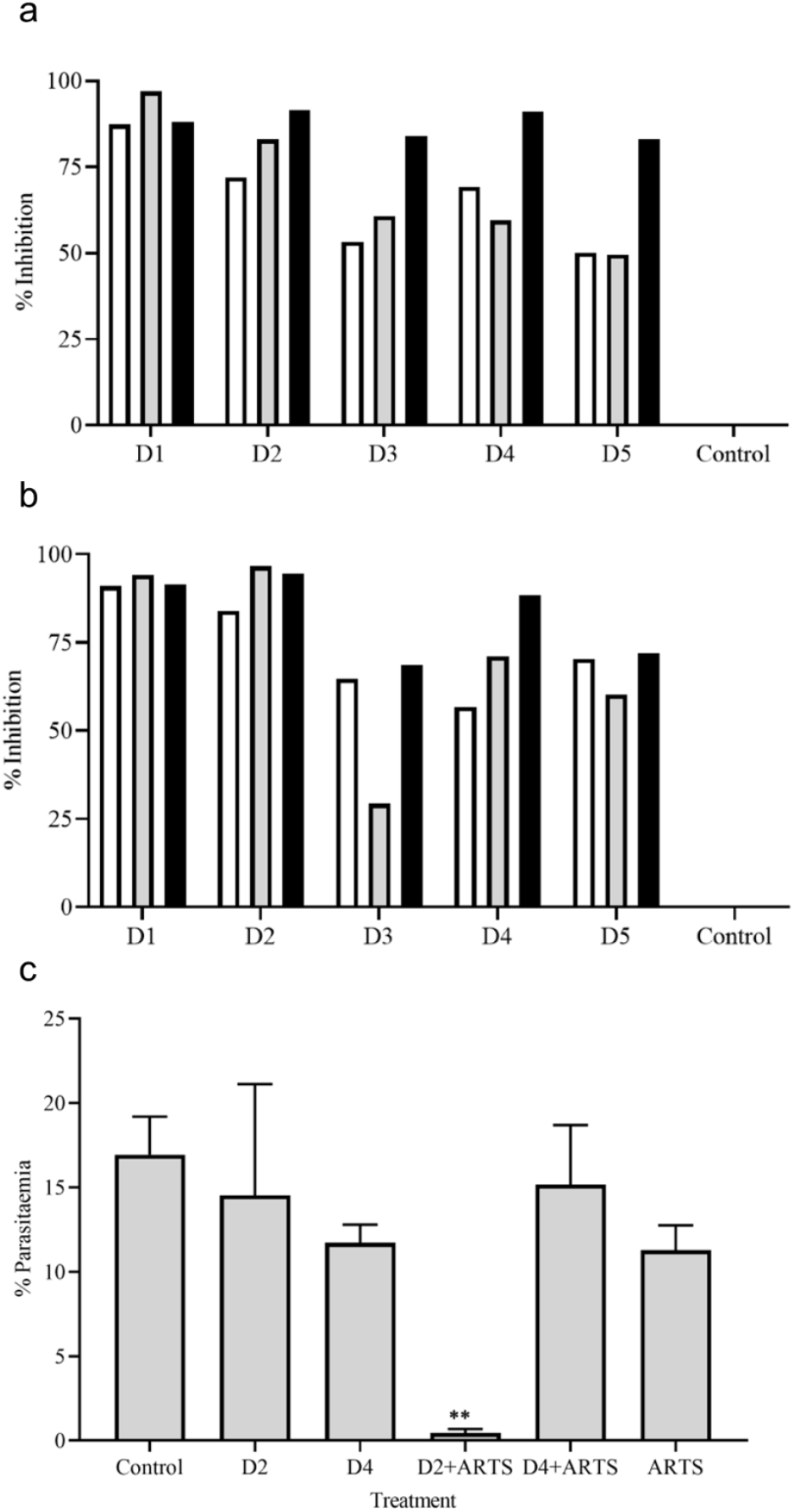
Anti-plasmodial activity of *V. odorata* plant extracts. The percent growth inhibition of 3D7 (□), RKL -9 (■) and R539T (■) strains at 48 (a) and 72 (b) hours. D1 - natural plant biomass extract, D2 - *In vitro* plant biomass extract, D3 - somatic embryo biomass extract, D4 -stirrer tank reactor cultivated cell suspension biomass extract and D5 - Airlift reactor cultivated cell suspension biomass extract. The concentration of plant extract - 250 μg mL^-1^. The initial parasitemia was 1 % and 2 % HC maintained. The plot (c) *in vivo* anti-plasmodial activity of *V. odorata* plant extracts. The percent parasitaemia was calculated from the thin tail blood smears stained with Giemsa. The sample was collected at 4^th^ day of post-infection. The concentration of plant extract was 400 mg kg^-1^ d^-1^, and ARTS was 1.8 mg kg^-1^ d^-1^. ARTS -artesunate. Different plant biomass extracts and their combination with ARTS were compared with the control. **p <0.01.

The natural plant extracts alone did not show any growth inhibition against *P. berghei* (Fig. 8c). The plant extracts only when combined with the standard anti-malarial drug ARTS showed parasite growth inhibition (Fig. 8c). The *in vitro* plant extracts (400 mg kg^-1^ d^-1^) combined with ATRS (1.8 mg kg^-1^ d^-1^) showed significant inhibition in the mice model, and the microscopic observations are presented in Fig. 9a, and b. Malaria infection could alter vascular permeability within treated mice’s blood–brain barrier (BBB). The blue staining of Evans blue dye in control mice, in contrast to D2+ARTS treated mice, indicated that the extract did not induce any observable changes in the BBB and exhibited protective effects against malarial parasites. Mice brains treated with Evans blue showed the compromised (Fig. 9c) and uncompromised (Fig. 9d) blood-brain barrier

**Fig 9.**
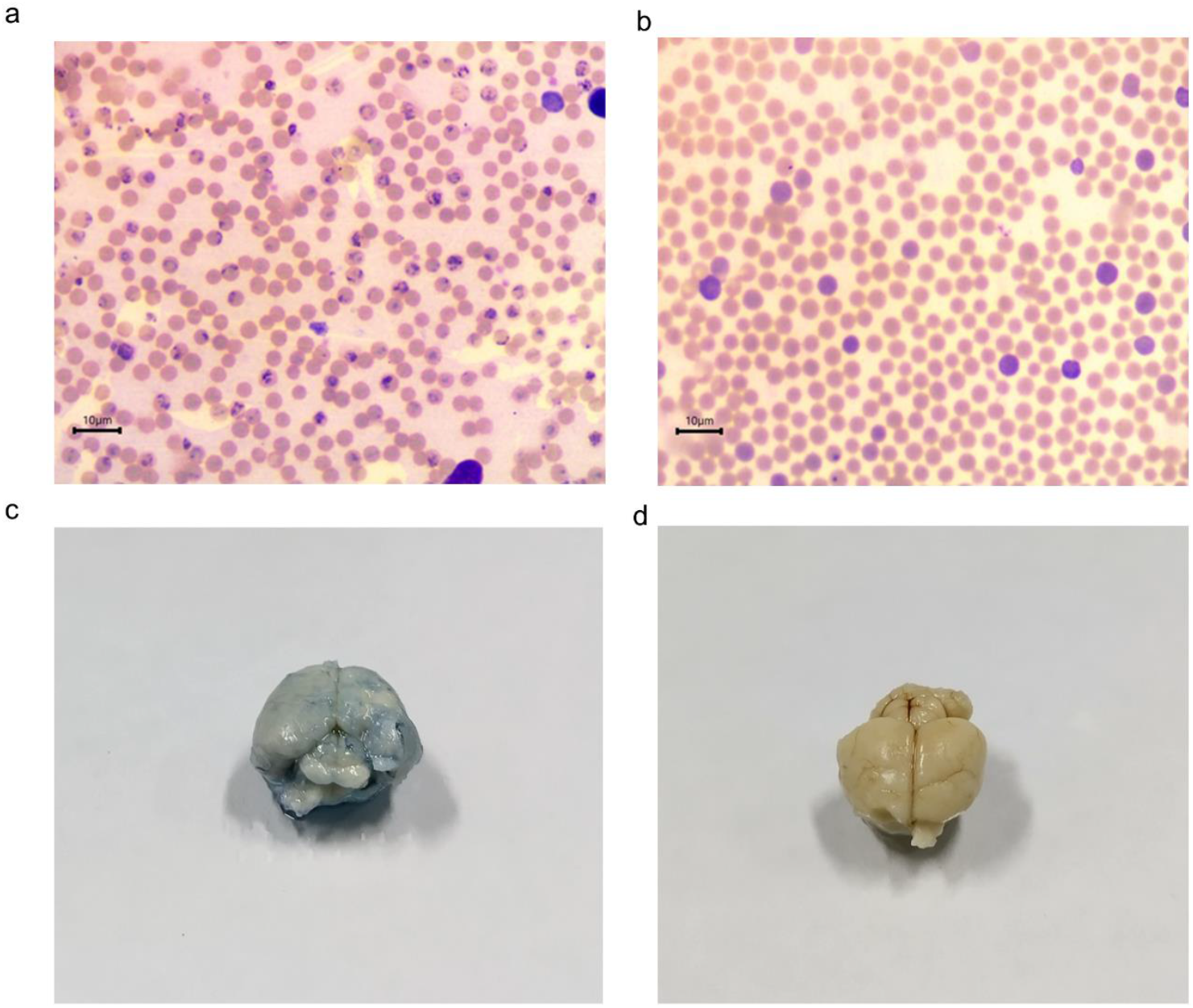
Anti-plasmodial activity of *V. odorata* plant extracts. Giemsa staining of thin blood smear as 100x magnification, and the arrows indicate the presence of parasites on the 4^th^ day. The figure (a) control and (b) D2+ARTS combination. D2- *in vitro* plant extract and ARTS -artesunate. The concentration of plant extract was 400 mg kg^-1^ d^-1^, and ARTS was 1.8 mg kg^-1^ d^-1^. Figures (c) and (d) showed the Evans blue staining of control and D2+ARTS treated mice’s brains.

Cyclotides were discovered from *O. affinis*, a medicinal plant found in African regions. In sub-Saharan Africa, people used *O. affinis* to treat malarial fever. The antiplasmodial activity of *O. affinis* has recently been reported (Nworu et al. 2017). The present study evaluated the efficacy of *V. odorata* natural and *in vitro* generated plant biomass extract (cyclotide-rich extract) against *Plasmodium* parasites in *in vitro* (at 250 μg mL^-1^) and *in vivo* (at 400 mg kg^-1^d^-1^) conditions and showed the parasite growth inhibition. In *in vivo* studies, a plant extract obtained from *O. affinis* inhibited *Plasmodium berghei* growth with a dose of 400 mg kg^-1^d^-1^. Also, the plant extract showed no harmful effects up to 5000 mg kg^-1^ with oral administration (Nworu et al. 2017). These results are aligned with the current experimental results, as *V. odorata* and *O. affinis* plant extracts are rich in cyclotides. Recently, Mann et al. (2023) reported the antiplasmodial activity of a cyclotide-rich plant extract obtained from *V. canescens* Wall against *P. falciparum* strain Pf 3D7. The results showed promising activity of these extracts, suggesting their potential as effective anti-malarial agents. As demonstrated in the current study, plant extracts can be combined with standard anti-malarial drugs to control parasite growth.

## 4. Conclusion

In this study, the bioreactor cultivation protocol for *V. odorata* cell suspension culture was developed. Modified STR and BTBCB were used to cultivate *V. odorata* cell suspension. The shake flask optimised conditions were implemented in the bioreactor, and the shake flask biomass productivity (1.8 g DW L^-1^ d^-1^) was replicated in the 3 L bioreactors. Further, the batch kinetic models were developed, and the model predictions were validated with experiments. A combination of the Monod and Sigmoid model was extrapolated into the fed-batch model to develop a suitable nutrient feeding strategy. The selected feeding strategy resulted in a biomass concentration of 32.2 g DW L^-1^ with a modified STR. However, the overall biomass productivity did not increase significantly; the biomass concentration increased 3-fold compared to the unoptimized control conditions (10.2 ± 0.8 g DW L^-1^). The plant extract obtained from the bioreactor-cultivated biomass was tested for cytotoxic, hemolytic, anti-inflammatory, and anti-plasmodial activities. In particular, the biomass extracts obtained from STR (D4) showed promising cytotoxic activity (IC50: 1.5 ± 0.1 mg mL^-1^) against Caco-2 human colorectal adenocarcinoma cells. In addition, the *in vitro*-generated biomass extracts inhibited parasite growth by up to 80 % in all three tested strains. In the case of *in vivo* studies, the *in vitro-generated* biomass extract (D2) combined with lower concentrations of ARTS (1.8 mg kg^-1^d^-1^) inhibited parasite growth. From the metabolite analysis and bioactivity assays, it can be said that the *in vitro-generated* plant biomass has similar medicinal potential to that in the natural plant source. Hence, *V. odorata* biomass generated using plant cell bioprocessing methods can be proposed an alternative to natural plant biomass for its medicinal and commercial applications.

Mass transfer limitations, excessive foaming and cell adhesion on the bioreactor walls are the major hindrances to *V. odorata* biomass production in high-cell-density cultivation. To overcome these limitations, computational tools such as Computational Fluid Dynamics (CFD) can be used to understand system requirements and design a plant cell-specific bioreactor configuration. Single-use bioreactor bags designed with non-sticky materials could avoid *V. odorata* cell attachment on bioreactor walls. Chemostat and turbidostat tools could be implemented to enhance *V. odorata* biomass productivity. For continuous and sustainable production, optimization of the cryopreservation protocol is required to facilitate long-term storage and scale-up into a pilot-scale bioreactor that supports the commercial-scale cultivation of *V. odorata* cell suspension.

## Acknowledgments

Prof. Smita Srivastava would like to acknowledge the financial support under Corporate Social Responsibility (CSR) by Hyclone Life Sciences Solutions India Private Limited (Cytiva) (Project Number: CR22230114BTHLSS008458) and L&T Technology Services (Project Number: CR21221810BTLNTE008458) for the ongoing research in the plant cell technology lab at Indian Institute of Technology Madras including the work stated above.

Babu R would like to acknowledge the Science & Engineering Research Board (SERB), Department of Science and Technology (DST), Government of India, Confederation of Indian Industry (CII) and Himalaya Wellness Company for the Prime Minister’s Fellowship Scheme for Doctoral Research (SERB/PM Fellow/CII-FICCI/Meeting/2019).

## Author contributions

BR: Methodology, Investigation, Data curation, Formal analysis, Visualization, Validation, Writing – Original Draft. MV: Software, Methodology, Investigation, Visualization, AW: Methodology, Investigation, Visualization VM: Supervision for anticancer and hemolytic activity, Funding acquisition, Resources, Conceptualization, Formal analysis, SS: Supervision for anti-malarial and anti-inflammatory activity, Funding acquisition, Resources, Conceptualization, Formal analysis, SSV: Supervision for overall research work as principal investigator, Funding acquisition, Resources, Ideation, Conceptualization, Formal analysis, Validation, Project administration, Writing – Review & Editing.

(SSV: Smita Srivastava; SS: Shailja Singh; BR: Babu R; MV: Manokaran Veeramani; AW: Aadinath Wallepure; VM: Vignesh Muthuvijayan)

