## Supplementary file for "Model-based fed-batch cultivation of *Viola odorata* plant cells exhibiting antimalarial and anticancer activity"

**Model-based fed-batch cultivation of *Viola odorata* plant cells exhibiting antimalarial and anticancer activity**
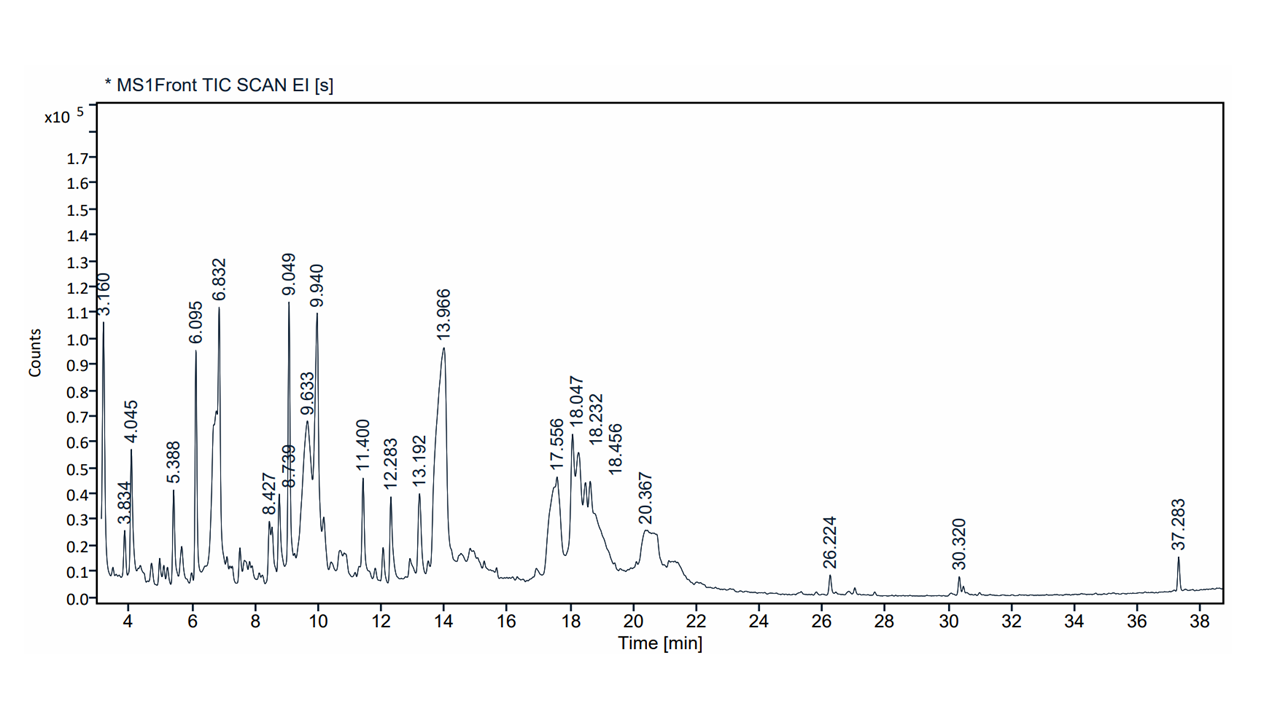

**Fig S1:** Full scan GCMS chromatograms of *V. odorata* biomass extracts obtained from STR operated in fed batch mode of cultivation.

**Table S1: Primary and secondary metabolites identified from *V. odorata* biomass extracts obtained from STR operated in fed batch mode of cultivation using GCMS**

| **S. No** | **RT** | **Compound name** |
| --- | --- | --- |
| 1. | 3.160 | 2-Cyclopenten-1-one, 2- hydroxy- |
| 2. | 3.834 | 2,4-Dihydroxy-2,5- dimethyl-3(2H)-furan-3- one |
| 3. | 4.045 | 2(5H)-Furanone, 5-ethyl- |
| 4. | 5.388 | 1,3,4,6,7,8-Hexahydro-1- methyl-2H-pyrimido[1,2- a]pyrimidine |
| 5. | 6.095 | cis-Aconitic acid |
| 6. | 6.832 | 4H-Pyran-4-one, 2,3- dihydro-3,5-dihydroxy-6- methyl- |
| 7. | 8.427 | 1,2,6-Hexanetriol |
| 8. | 8.739 | 5-Hydroxymethylfurfural |
| 9. | 9.049 | L-Sorbose |
| 10. | 9.633 | D-Galactose |
| 11. | 9.940 | 5-(2-Hydroxyethyl)-4- methylthiazole |
| 12. | 11.400 | Isosorbide Dinitrate |
| 13. | 12.283 | 1H-Pyrazole-3-acetic acid, 2,5-dihydro-5-oxo |
| 14. | 13.192 | Butanamide, 2-hydroxy-N,2,3,3-tetramethyl- |
| **S. No** | **RT** | **Compound name** |
| 15. | 13.966 | Sucrose |
| 16. | 17.556 | 3-Deoxy-d-mannoic lactone |
| 17. | 18.047 | Trehalose |
| 18. | 18.232 | d-Glycero-d-ido-heptose |
| 19. | 18.456 | d-Mannose |
| 20. | 20.367 | β-D-Glucopyranose |
| 21. | 26.224 | n-Hexadecanoic acid |
| 22. | 30.320 | 1,E-11,Z-13- Octadecatriene |
| 23. | 37.283 | Hexadecanoic acid, 2- hydroxy-1- |
